# AZIN2-dependent polyamine metabolism determines adipocyte progenitor fate and protects against obesity and dysmetabolism

**DOI:** 10.1101/2024.11.19.621837

**Authors:** Christine Mund, Anupam Sinha, Anika Aderhold, Ivona Mateska, Eman Hagag, Sofia Traikov, Bettina Gercken, Andres Soto, Jonathan Pollock, Lilli Arndt, Michele Wölk, Natalie Werner, Georgia Fodelianaki, Pallavi Subramanian, Kyoung-Jin Chung, Sylvia Grossklaus, Mathias Langner, Mohamed Elgendy, Tatyana Grinenko, Ben Wielockx, Andreas Dahl, Martin Gericke, Matthias Blüher, Ünal Coskun, David Voehringer, Maria Fedorova, Mirko Peitzsch, Peter J. Murray, Triantafyllos Chavakis, Vasileia Ismini Alexaki

## Abstract

Adipose tissue homeostasis plays a critical role in metabolic disease but the metabolic circuitry regulating adipose tissue dynamics remains unclear. In this study, polyamine metabolism emerges as an important regulator of adipose tissue pathophysiology. We identify AZIN2 (Antizyme inhibitor 2), a protein promoting polyamine synthesis and acetylation, as a major regulator of total acetyl-CoA in adipocyte progenitors (APs). AZIN2 deficient APs demonstrate increased H3K27 acetylation marks in genes related to lipid metabolism, cell cycle arrest and cellular senescence, and enhanced adipogenesis compared to wild-type counterparts. Upon high-fat diet (HFD)-induced obesity, global AZIN2 deficiency in mice provokes adipose tissue hypertrophy, AP senescence, lipid storage perturbations, inflammation and insulin resistance. IL4 promotes *Azin2* expression in APs but not mature adipocytes due to diminished IL4 receptor expression in the latter. In human visceral and subcutaneous adipose tissue, *AZIN2* expression positively correlates with expression of early progenitor markers and genes associated with protection against insulin resistance, while it negatively correlates with markers of lipogenesis. In sum, AZIN2-driven polyamine metabolism preserves adipose tissue health, a finding that could be therapeutically harnessed for the management of obesity-associated metabolic disease.

## Introduction

Obesity promotes development of type 2 diabetes, fatty liver disease, and cardiovascular disease ^1^. Although the ability of adipocytes to sequester fat protects other organs from lipotoxicity ^2^, the excessive expansion of adipose tissue, leading to hypoxia and cell destruction, is associated with development of chronic low-grade inflammation and whole body dysmetabolism ^3, 4^. Adipose tissue growth is driven by the increase in size (hypertrophy) and number (hyperplasia) of adipocytes ^2^. In obesity, adipose tissue hyperplasia is linked to metabolic advantages compared to adipose tissue hypertrophy ^4^. On the other hand, generation of new adipocytes (adipogenesis) from adipocyte progenitors (APs) is triggered at the start of obesity facilitating adipose tissue expansion ^5–8^.

New adipocytes emerge from platelet-derived growth factor-α positive (PDGFRα^+^) progenitors ^9, 10^. Single-cell transcriptomic studies revealed different subpopulations of APs in human and mouse adipose tissue with variable adipogenic capacity ^11–15^. The molecular path to de novo adipogenesis from APs is controlled in part by CCAAT-enhancer-binding protein β (C/EBPβ), C/EBPδ, and Peroxisome Proliferator-Activated Receptor γ (PPARγ), leading to accumulation of lipid droplets that gradually merge to form mature adipocytes ^4, 16^, which further increase in size by triglyceride (TG) storage. Although the transcriptional regulation of adipogenesis has been deciphered in detail ^16^, its regulation by metabolic pathways remains largely unknown. Polyamines are present in all life kingdoms ^17^. As polycations, they bind to negatively charged molecules, such as DNA, RNA or phospholipids, and have broad functions in cellular homeostasis and division ^18–23^. Polyamine amounts are regulated by de novo biosynthesis (from arginine or glutamine), recycling and uptake by transporters ^24, 25^. Arginine is metabolized through arginase (ARG) 1 or 2 to L-ornithine, which is further metabolized by ornithine decarboxylase 1 (ODC1) to putrescine, the latter being the rate-limiting step of de novo polyamine biosynthesis. Putrescine is converted to spermidine by spermidine synthase (SRM), and spermidine is used by spermine synthase (SMS) to produce spermine ^23^. In recycling reactions, spermine can regenerate spermidine via spermine oxidase (SMOX). Spermidine is acetylated by spermidine/spermine-N1-acetyltransferase (SAT1) to N1-acetyl-spermidine, which is oxidized by polyamine oxidase (PAOX) to regenerate putrescine ^26^. Polyamine synthesis is primarily regulated at the level of ODC1 by antizymes (AZ1-3), which physically interact with ODC1 to inhibit its activity and stimulating its ubiquitin-independent proteasomal degradation ^21, 24^. In turn, AZs are themselves regulated by two antizyme inhibitors (AZIN1 and 2), which are structurally similar to ODC1 but enzymatically inactive and titrate AZs, freeing ODC1 to form active homodimers ^27, 28^. Thus, AZIN1 and AZIN2 by virtue of their ability to inhibit the inhibitors, are positive regulators of ODC1.

We demonstrate here that polyamine metabolism plays a profound role in adipose tissue homeostasis. Numerous components of polyamine metabolism are expressed in APs and their expression in adipose tissue is downregulated in obesity. Amongst them, we found that AZIN2 is a key regulator of adipogenesis. AZIN2 promotes polyamine synthesis and acetylation, decreases acetyl-CoA amounts and negatively regulates H3K27 and H3K9 acetylation marks in regulatory elements of adipogenic genes in APs. Young chow diet (CD)-fed *Azin2^-/-^* mice have increased adipogenesis and reduced AP proliferation leading to diminished AP numbers. Ηigh-fat diet (HFD)-fed *Azin2^-/-^*mice develop enlarged adipose tissue with more prominent inflammation and AP senescence, impaired insulin sensitivity, reduced energy expenditure and enhanced liver steatosis compared to wild-type (wt) mice indicating that AZIN2-driven polyamine metabolism in APs is critical for maintaining adipose tissue health. In sum, AZIN2-driven polyamine metabolism emerges as a potent regulator of adipose tissue homeostasis.

## Results

### Polyamine metabolism declines with obesity

We analyzed the inguinal subcutaneous adipose tissue (SAT) of mice fed for 20 weeks a low- or a high-fat diet (LFD and HFD, respectively) by bulk RNA-seq. HFD feeding induced body weight gain, glucose intolerance, and insulin resistance ^29, 30^ and caused transcriptional remodeling of the adipose tissue (Figure S1A). Expression of adipogenesis- and lipid metabolism-related genes was upregulated in the adipose tissue of HFD mice (Figure S1B). Furthermore, we observed enrichment of inflammatory response-related changes (Figure S1C,D) in the adipose tissue of obese mice in accordance with the well-documented pro-inflammatory profile of the adipose tissue in obesity ^3^. By contrast, mRNAs encoding proteins mediating arginine and proline metabolism were downregulated in the adipose tissue of HFD mice (Figure 1A-C, S1E), among which was *Azin2*, *Odc1* and *Sms* (Figure 1B-D), suggesting that HFD downregulated polyamine metabolism in the adipose tissue.

**Figure 1.**
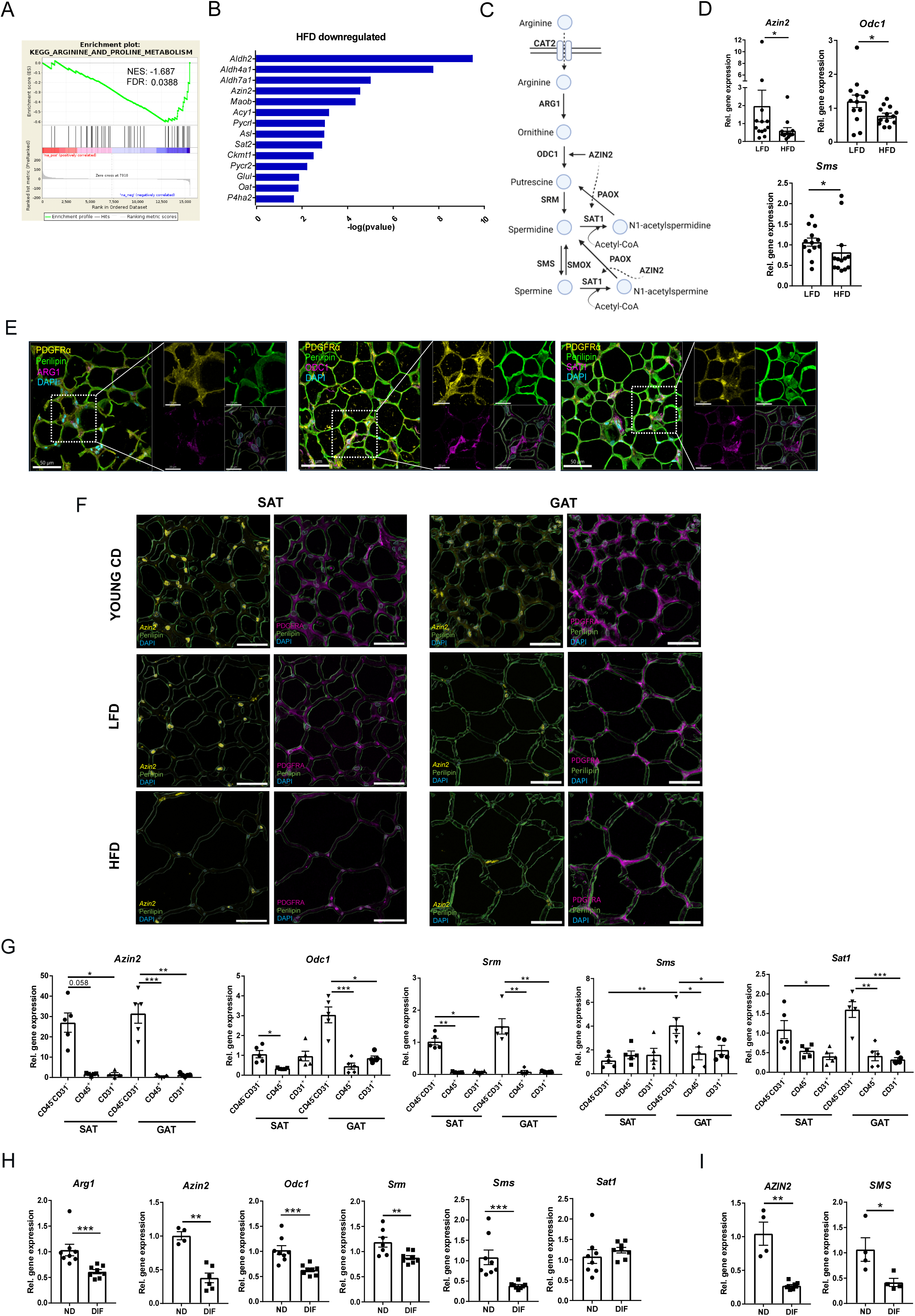
Expression of proteins involved in polyamine metabolism is downregulated in adipose tissue with obesity. **A.** GSEA analysis of bulk RNA-seq in SAT of mice fed for 20 weeks a HFD or LFD for KEGG_arginine and proline metabolism (n=4 mice per group). **B.** Arginine metabolism-related genes downregulated in SAT of HFD mice based on RNA-seq data (n=4 mice per group). **C.** Schematic presentation of polyamine metabolism. **D.** *Azin2*, *Odc1* and *Sms* relative gene expression in SAT of mice fed for 20 weeks a HFD or LFD (n=13 mice per group). **E.** Immunofluorescence for ARG1 (magenta), ODC1 (magenta) and SAT1 (magenta), PDGFRα (yellow), Perilipin (green) and DAPI staining in SAT of mice fed a CD. Scale bar, 50 μm. Representative images are shown. **F.** RNAScope for *Azin2* and immunofluorescence for PDGFRα and perilipin in SAT and GAT of 8 week-old mice fed a CD and mice fed for 20 weeks a LFD or a HFD. Scale bar, 50 μm. Representative images from 2 mice, 3 levels per tissue are shown. **G.** *Azin2*, *Odc1*, *Srm*, *Sms* and *Sat1* expression in CD45^+^, CD31^+^ and CD45^-^CD31^-^ SVF populations isolated from SAT and GAT of wt mice (n=5 mice per group). **H.** *Arg1*, *Azin2*, *Odc1*, *Srm*, *Sms* and *Sat1* expression in differentiated (DIF) or non-differentiated (ND) cultured murine SAT APs (n=4-8 mice per group). **I.** *AZIN2* and *SMS* expression in differentiated or non-differentiated cultured human preadipocytes (n=4-8). Gene expression in **D,G-I** was determined by qPCR using *18S* as a housekeeping gene. Data in **D,G-I** are shown as mean±SEM. Mann-Whitney U (**D,H,I**) or Kruskal-Wallis (**G**) tests were used for statistical analysis. *p < 0.05, **p < 0.01, ***p < 0.001. NES: Normalized enrichment score; FDR: False discovery rate

Next, we set out to identify cells of the adipose tissue expressing components of polyamine metabolism. At the protein level, we found that ARG1, ODC1 and SAT1 are expressed in PDGFRα^+^ APs in adipose tissue of 8 week-old CD-fed C57BL/6J mice (Figure 1E). Because we do not have a specific antibody for AZIN2, we used RNAscope in PDGFRα^+^ APs in SAT and gonadal adipose tissue (GAT) (Figure 1F, S1F). *Pdgfra* was expressed in CD45^-^CD31^-^ cells of the stromal-vascular fraction (SVF) (Figure S1G) and was not expressed in the adipocyte fraction (AF) consisting of mature adipocytes (Figure S1H). *Azin2*, *Odc1*, *Srm*, *Sms* and *Sat1* expression was detected in CD45^-^CD31^-^ SVF cells and their expression was higher in CD45^-^CD31^-^ compared to CD45^+^ and CD31^+^ SVF cells (Figure 1G). *Arg1* expression was similar in CD45^-^ CD31^-^, CD45^+^, CD31^+^ SVF cells (data not shown). Although *Azin2* mRNA was detected at relatively low amounts in the white adipose tissue compared to other tissues, such as testes (not shown), the brain or the adrenal glands (Figure S1I), *Azin2* was highly expressed in APs compared to other cell types, such as mature adipocytes, adipose tissue macrophages (ATMs), neutrophils and hepatocytes (Figure S1J).

*Azin2* mRNA was diminished in the adipose tissue of mice fed for 20 weeks a HFD compared to young CD-fed mice or mice fed for 20 weeks a control LFD (Figure 1F). This was mainly due to proportional reduction of AP numbers and low *Azin2* mRNA expression in mature adipocytes (Figure 1F, S1J, S1K). In accordance, the expression of *Azin2*, *Arg1*, *Odc1, Srm* and *Sms* declined upon in vitro differentiation of mouse APs towards adipocytes (Figure 1H, S1L), even though full adipocyte maturity (characterized by appearance of a single large lipid droplet) was not reached in this in vitro system, in which multilocular adipocytes formed instead (Figure S4H,J). Similarly, *AZIN2* and *SMS* mRNA expression was also diminished upon differentiation in human preadipocytes (Figure 1I). Thus, key components of polyamine metabolism, including AZIN2, are expressed in APs and their expression declines with adipocyte differentiation.

### IL4 upregulates *Azin2* expression in APs

Next, we set out to investigate whether factors deriving from the adipose tissue microenvironment regulate polyamine metabolism in APs. Given that in the adipose tissue polyamine metabolism is downregulated (Figure 1A,B,D,F) and the immune profile radically changes with obesity (Figure S1C,D) ^3, 31^, we considered immune signals may regulate polyamine metabolism in APs. IL4 is a type 2 immunity cytokine, which targets preadipocytes ^32^. In adipose tissue, eosinophils, sustained by IL33 and IL5-driven signaling networks, are a major source of IL4 ^33, 34^. IL4Rα is expressed in SAT APs (Figure S2A), and its expression is reduced upon adipogenesis (Figure 2A). Accordingly, *Il4ra* expression is higher in CD11b^-^ SVF cells than in the AF (Figure S2B). IL4 induced transcriptional reprograming of cultured non-differentiated APs (sorted as PDGFRα^+^LY6A^+^CD31^-^CD45^-^ cells) (Figure 2B). Arginine and proline metabolism was the top amongst the metabolic pathways with positive enrichment as shown by EGSEA analysis (Figure 2C) and *Azin2*, *Arg1* and *Srm* were among the upregulated arginine metabolism-related mRNAs (Figure 2D). IL4 also increased the expression of different amino acid transporters of the solute carrier family 7 (SLC7), among which *Slc7a2*, encoding for the arginine transporter CAT2 (cationic amino acid transporter 2) was one of the most strongly upregulated (Figure 2E,F) ^35^. Similarly, IL4 increased *AZIN2* mRNA expression in human preadipocytes (Figure 2G). The effect of IL4 on *Azin2* expression was abolished by silencing of STAT6, the transcription factor mediating the majority of the effects of IL4 and IL13 (Figure S2C) ^36^. IL13, which exerts its effects through the type II receptor IL4Rα/IL13Rα1 in non-hematopoietic cells ^37^, also increased *Azin2* expression in mouse APs (Figure S2D). Moreover, IL4 and IL13 upregulated *Azin2*, *Slc7a2* and *Arg1* expression in adipose tissue explants from wild type (wt) but not *Il4ra^-/-^*mice (Figure 2H).

**Figure 2.**
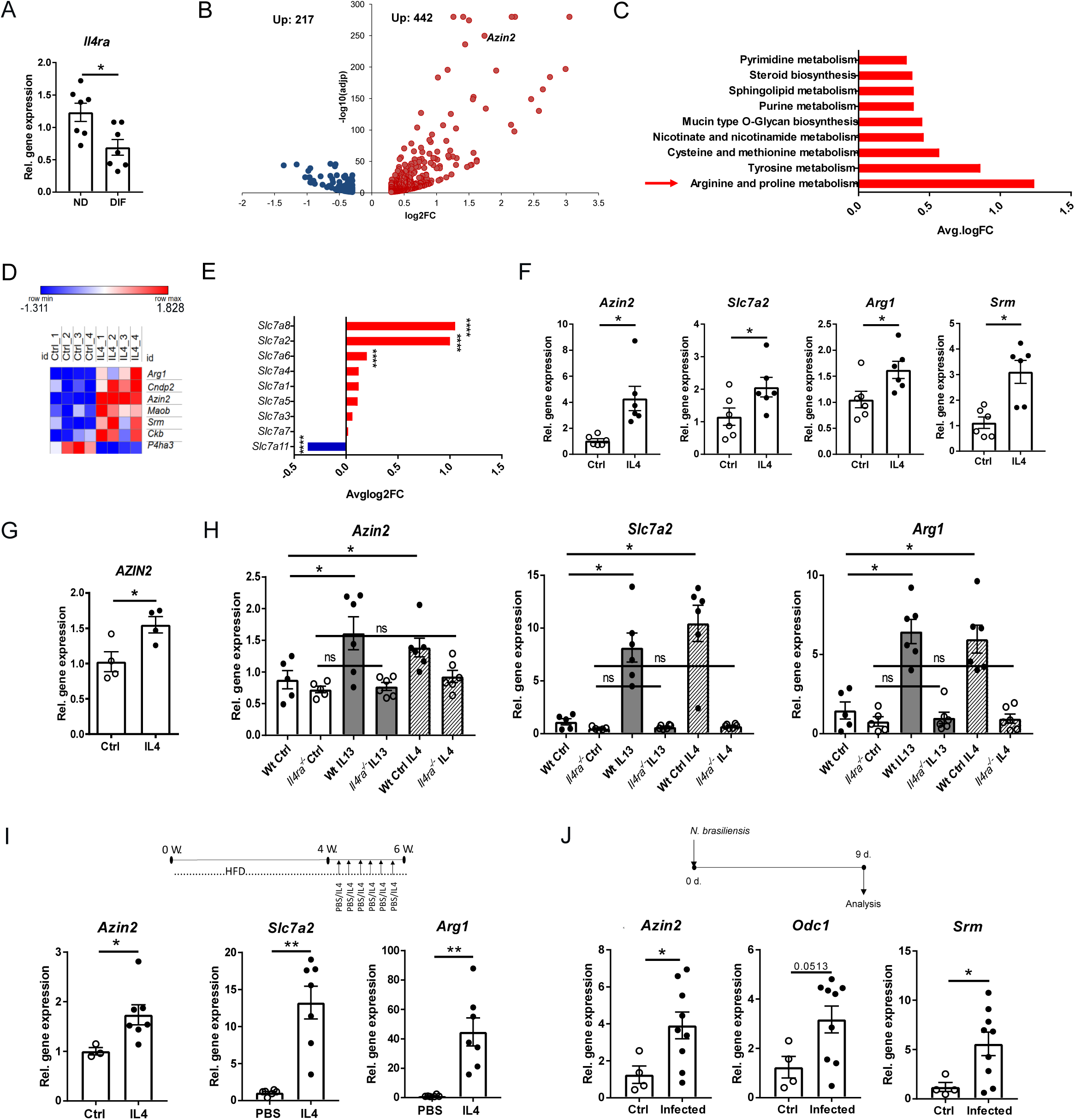
IL4 upregulates *Azin2* expression in APs. **A.** *IL4ra* expression in differentiated and non-differentiated SAT APs assessed by qPCR (n=7 mice per group). **B-E.** Bulk RNA-seq was performed in mouse SAT APs treated for 4 h with IL4 (20 ng/ml). Volcano plot for differentially expressed genes (DEG) with padj<0.05 log2FC>0.3 and < -0.3 (**B**), positively enriched KEGG metabolic pathways based on EGSEA analysis (**C**), heatmap for arginine metabolism genes (**D**), expression of Slc family 7 cationic amino acid transporters (**E**) (n=4 mice per group for **B-E**). **F.** *Azin2, Slc7a2, Arg1* and *Srm* expression in mouse APs treated for 4 h with IL4 (20 ng/ml) (n=6 mice per group). **G.** *AZIN2* expression in human preadipocytes treated for 24 h with IL4 (12 ng/ml) (n=4). **H.** *Azin2, Slc7a2* and *Arg1* expression in GAT explants from *Il4ra^-/-^* and wt mice treated for 4 days with 50 ng/ml IL4 or IL13 (n=5-6 mice per group). **I.** *Azin2* expression in SAT SVF, and *Slc7a2* and *Arg1* in SAT of mice fed for 6 weeks a HFD and treated every 2 days with PBS or IL4 + a-IL4 during the last 2 feeding weeks, as depicted in the scheme (n=3-7 mice per group). **J.** *Azin2, Odc1* and *Srm* expression in GAT of mice infected with *N. brasiliensis* or control mice 9 days post-infection (n=4-9 mice per group). Gene expression in **A,F-J** was determined by qPCR using *18S* as a housekeeping gene and shown as mean±SEM. Mann-Whitney U test was used for statistical analysis in **A**,**G,I,J**; Wilcoxon test was used for statistical analysis in **F**, Kruskal-Wallis test was used for statistical analysis in **H** *p < 0.05, **p < 0.01.

IL4 was previously reported to inhibit adipogenesis ^38, 39^. In accordance, wt mice treated with IL4 during the 2 last weeks of a 6-week-long HFD feeding showed reduced body weight gain (Figure S2E), diminished expression of adipogenic genes, such as *Pparg*, CCAAT/enhancer-binding protein-alpha (*Cebpa*), Acyl-CoA Synthetase Long Chain Family Member 1 (*Acsl1*), Adiponectin (*Adipoq*), *Cd36* and lipase, hormone sensitive (*Lipe*) (Figure S2F) and increased AP number per AT weight (Figure S2G). As expected, expression of genes encoding for proteins related to type 2 immunity, such as *Arg1*, chitinase-like 3 (*Chil3*), *resistin like alpha (Retnla)*, *C-type lectin domain family 10* (*Clec10a*), and *mannose receptor, C type 1* (*Mrc1*), was increased in the SAT SVF of IL4-compared to PBS treated mice (Figure S2H) ^36^. In accordance, IL4 increased *Azin2*, *Slc7a2* and *Arg1* expression in SAT SVF of mice (Figure 2I).

Furthermore, infection of mice with *N. brasiliensis*, which causes a potent type 2 immune response and eosinophil accumulation in the adipose tissue ^40, 34^, also downregulated *Pparg* and *Cd36* expression (Figure S2I) and increased *Azin2* and *Srm* expression in GAT (Figure 2J).

In order to validate these findings in human adipose tissue, we mined the public GEPIA2 database (http://gepia2.cancer-pku.cn/#index) ^41^ for the expression of type 2 immune mediators in correlation with *AZIN2* expression. Expression of *IL4, IL13, STAT6, IL5* and *IL33* positively correlated with *AZIN2* expression in subcutaneous and visceral adipose tissue (Figure S2J-R). These data collectively suggest that type 2 immunity upregulates *Azin2* expression in APs.

### AZIN2 promotes polyamine acetylation and regulates acetyl-CoA amounts in APs

We next generated mice with global *Azin2* deletion. *Azin2* expression was abolished in SAT APs of *Azin2^-/-^* mice (Figure S3A). GAT-derived *Azin2^-/-^* APs, cultured in a spermidine-free medium in order to exclude AZIN2-dependent differences in spermidine uptake ^42^, had lower intracellular spermidine amounts compared to wt APs (Figure 3A left panel), thereby validating that AZIN2 promotes ODC1 function and spermidine production ^21^.

**Figure 3.**
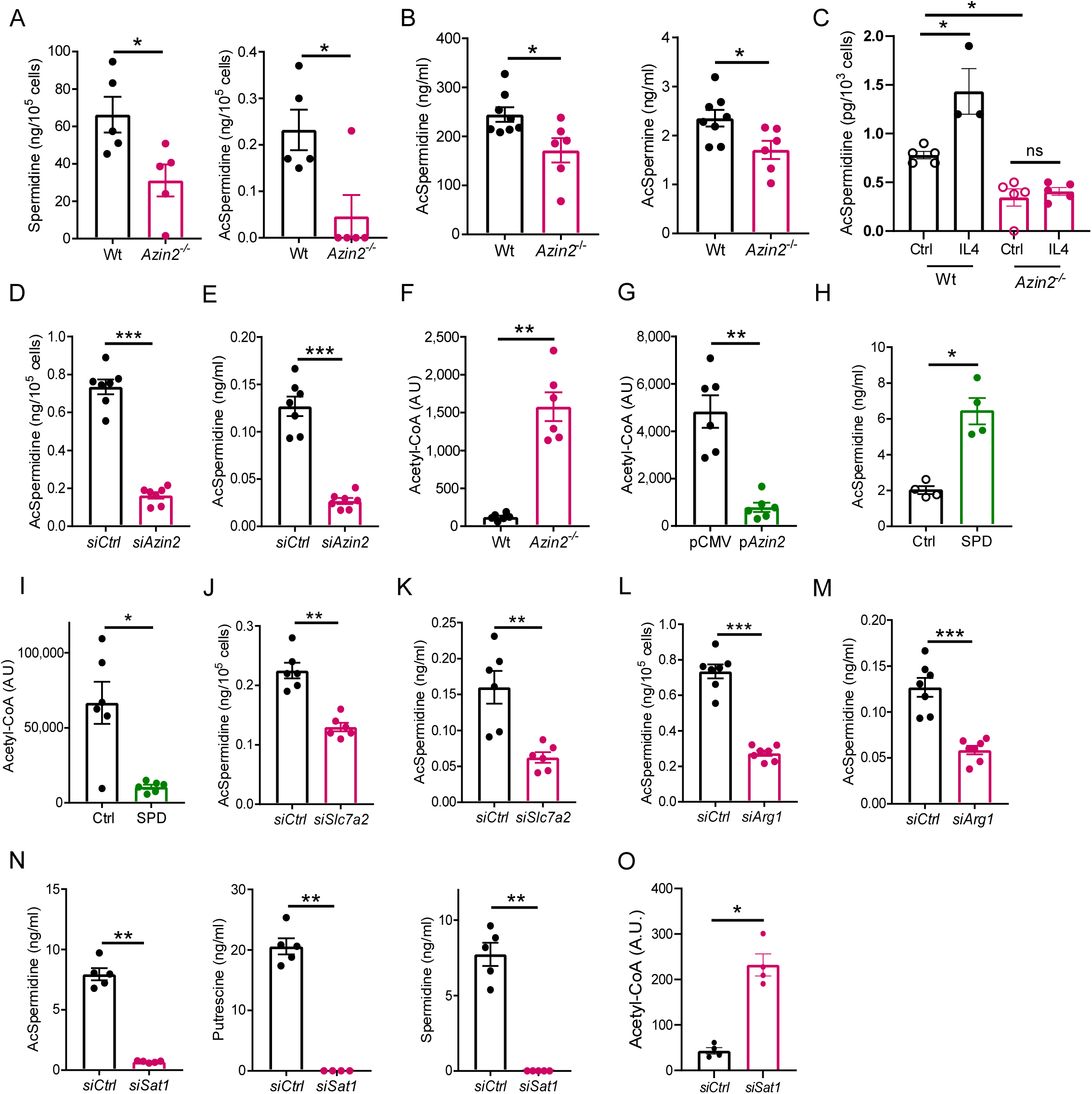
AZIN2 promotes polyamine acetylation and negatively regulates acetyl-CoA levels in APs. **A.** Intracellular spermidine and N1-acetylspermidine levels in undifferentiated cultured wt and *Azin2^-/-^*GAT APs (n=5 mice per group). **B.** N1-acetylspermidine and N1-acetylspermine concentrations in supernatants of GAT explants from wt and *Azin2^-/-^* mice kept 18 hours in culture (n=6-8 mice per group). **C.** N1-acetylspermidine concentration in cell lysates of wt and *Azin2^-/-^* SAT APs treated for 24 h with IL4 (20 ng/ml) (n=3-5 mice per group). **D,E.** SAT APs were transfected 3 times over one week of differentiation with si*Azin2* or siCtrl and N1-acetylspermidine was measured in the cell lysates (**D**) and cell culture supernatants (**E**) (n=7 mice per group). **F.** Relative abundance of acetyl-CoA in non-differentiated wt and *Azin2^-/-^* SAT APs shown as peak intensity assessed by LC-MS/MS (n=6 mice per group). **G.** Relative abundance of acetyl-CoA levels in 3T3-L1 preadipocytes transfected with an *Azin2* overexpressing plasmid or a control plasmid (pCMV) shown as peak intensity assessed by LC-MS/MS (n=6 biological replicates). **H,I.** N1-acetylspermidine concentration in cell culture supernatants (**H**) and relative abundance of acetyl-CoA (**I**) in SAT APs treated with spermidine (SPD, 10 μΜ) during 1 week of differentiation (n=4 for **H** and 6 for **I** mice per group). **J,K.** N1-acetylspermidine concentrations in cell lysates (**J**) and cell culture supernatants (**K**) of undifferentiated SAT APs transfected with si*Slc7a2* or siCtrl (n=6 mice per group). **L,M.** N1-acetylspermidine concentrations in cell lysates (**L**) and cell culture supernatants (**M**) of SAT APs transfected 3 times over one week of differentiation with si*Arg1* or siCtrl (n=7 mice per group). **N.** N1-acetylspermidine, putrescine and spermidine concentrations in cell culture supernatants of undifferentiated SAT APs transfected with si*Sat1* or siCtrl (n=5 mice per group). **O.** Relative abundance of acetyl-CoA in non-differentiated APs transfected with si*Sat1* or siCtrl (n=3-4 mice per group). Data are shown as mean±SEM. Mann-Whitney U test was used for statistical analysis. *p < 0.05, **p < 0.01, ***p < 0.001. AU: Arbitary units. Data of ‘siCtrl’ samples are the same in **D** and **L**, and **E** and **M**.

Spermidine and spermine are modified by acetylation via SAT1 forming N1-acetylspermidine and N1-acetylspermine, respectively (Figure 1C). Intracellular N1-acetylspermidine amounts were diminished in GAT *Azin2^-/-^* APs (Figure 3A right panel). Accordingly, N1-acetylspermidine and N1-acetylspermine amounts were lower in supernatants of *Azin2^-/-^* GAT explants compared to supernatants of wt explants (Figure 3B). Moreover, N1-acetylspermidine amounts were reduced in culture supernatants of *Azin2^-/-^* SAT APs in comparison to wt counterparts (Figure S3B), suggesting that despite its reduced expression upon differentiation (Figure 1H), AZIN2 is still required for efficient polyamine acetylation in in vitro differentiated APs. Moreover, IL4 increased N1-acetylspermidine amounts in wt but not *Azin2^-/-^* SAT APs (Figure 3C), suggesting that IL4 promotes polyamine acetylation in an AZIN2-dependent manner. We also used *Azin2* siRNA silencing as a comparator loss-of-function approach to ensure that *Azin2^-/-^* mice-derived APs had not undergone substantial metabolic rewiring in terms of polyamine metabolism (Figure S3C). *Azin2* mRNA silencing reduced intracellular and exported amounts of N1-acetylspermidine in SAT APs (Figure 3D,E), decreased putrescine amounts and, almost significantly, increased ornithine levels (Figure S3D), confirming the promoting effect of AZIN2 on ODC-mediated conversion of ornithine to putrescine ^21, 43, 28^.

Spermidine and spermine acetylation requires acetyl-CoA (Figure 1C) ^23^. Given that AZIN2 promotes polyamine acetylation (Figure 3A, B, D, E, S3B), we tested if AZIN2 regulates acetyl-CoA amounts in APs. Strikingly, total acetyl-CoA was increased in both undifferentiated and differentiated *Azin2*^-/-^ APs compared to wt AP controls (Figure 3F, S3E). *Azin2* cDNA overexpression through plasmid transfection (Figure S3F) reduced acetyl-CoA in 3T3-L1 preadipocytes compared to pCMV-transfected cells (Figure 3G). Moreover, incubation of APs with spermidine fueled N1-acetylspermidine production (Figure 3H) and diminished acetyl-CoA amounts (Figure 3I), indicating that polyamine acetylation is driven by substrate availability and associates with reduction in acetyl-CoA amounts. These findings collectively suggest that AZIN2 promotes polyamine acetylation and is therefore a key regulator of acetyl-CoA amounts in APs. Next, we examined whether polyamine acetylation depends on arginine uptake and de novo polyamine synthesis. *Slc7a2* silencing (Figure S3G) reduced N1-acetylspermidine levels in SAT AP cell lysates and cell culture supernatants (Figure 3J,K). Similarly, *Arg1* silencing (Figure S3H) also reduced intracellular and exported N1-acetylspermidine levels in SAT APs (Figure 3L,M) suggesting that production of acetylated polyamines requires CAT2-dependent uptake of arginine and its conversion to ornithine and polyamines. Moreover, *Sat1* silencing (Figure S3I) blunted not only N1-acetylspermidine production but also putrescine and spermidine levels (Figure 3N), standing in accordance with the fact that N1-acetylspermidine regenerates putrescine through the polyamine flux (Figure 1C) ^23^. Furthermore, similarly to AZIN2 knockdown, *Sat1* silencing strongly increased acetyl-CoA amounts in APs (Figure 3O).

### AZIN2 inhibits de novo adipogenesis via regulation of histone acetylation

Acetyl-CoA is required for histone acetylation, a central mechanism of epigenetic regulation ^44^. As we found that AZIN2 downregulates acetyl-CoA levels (Figure 3G,H, S3E), we hypothesized that AZIN2 may indirectly affect histone acetylation in APs. H3K27ac demarks activated enhancers of key adipogenic genes ^45, 46, 16^. We performed ChIP-seq analysis for H3K27ac in in vitro differentiated *Azin2^-/-^* and wt SΑΤ APs and identified 10,953 sites of H3K27ac enrichment in *Azin2^-/-^*compared to wt APs (Figure 4A). These sites were detected in distal intergenic regions, promoter regions (2-3, 1-2 and < 1 kb), regions encoding for 5’ and 3’ UTR, exons and introns (Figure 4A, S4A). In wt APs 4,121 sites were H3K27ac enriched compared to *Azin2^-/-^* APs (Figure S4A). Accordingly, the genome-wide H3K27ac distribution of the ChIP-seq peaks across transcriptional start sites (TSS) (-/+ 5 kb) and in gene regions between TSS and transcription end sites (TES) was higher in *Azin2^-/-^* compared to wt APs (Figure 4B and S4B, respectively). Many sites with increased H3K27ac amounts in *Azin2^-/-^* APs were associated with genes encoding proteins linked to lipid metabolism, negative regulation of cell cycle, hypoxia and senescence (Figure 4C). A number of genes related to lipid metabolism showed greater H3K27ac enrichment in their promoter regions, exons, introns, regions encoding for 5’ UTR and distal intergenic regions in *Azin2^-/-^* compared to wt APs, such as *Pparg*, *Cebpa*, *Cebpd*, *Cebpe*, perilipin 4 (*Plin4*), stearoyl-coenzyme A desaturase 2 (*Scd2*), *Scd4*, acetyl-Coenzyme A acetyltransferase 2 (*Acat2*), acetyl-Coenzyme A carboxylase alpha (*Acaca*), ceramide synthase 3 (*Cers3*), lipid droplet associated hydrolase (*Ldah*), adiponectin receptor 2 (*Adipor2*), glycerol-3-phosphate acyltransferase 3 (*Gpat3*), peroxisome proliferative activated receptor, gamma, coactivator 1 alpha (*Ppargc1a*) and fatty acyl CoA reductase 1 (*Far1*) (Figure S4C-G).

**Figure 4.**
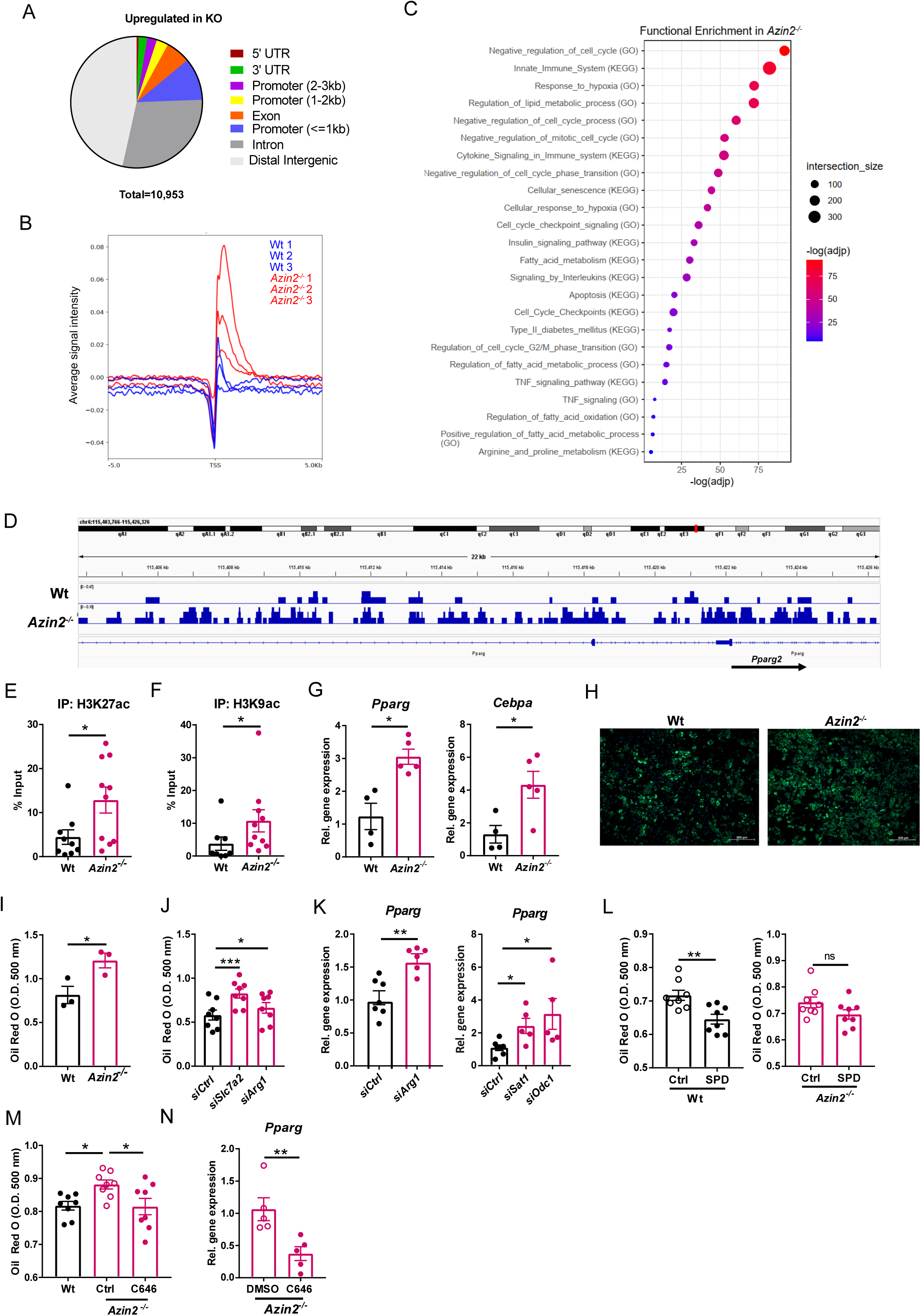
AZIN2 inhibits adipogenesis through regulation of histone acetylation. **A-E.** Bulk ChIP-Seq was performed after IP for H3K27ac in in vitro differentiated wt and *Azin2^-/-^*SAT APs (n=3 pools of APs from 2 mice per group). **A.** Global DNA distribution of H3K27ac marks upregulated in *Azin2^-/-^* SAT APs. **B.** Histogram of the H3K27ac signal intensity in wt and *Azin2^-/-^* SAT APs. **C.** Enriched GO terms and pathways and KEGG pathways in *Azin2^-/-^* compared to wt SAT APs plotted for the –log(adjp) value based on genes with enhanced H3K27ac signal in *Azin2^-/-^* versus wt APs. **D.** Integrative Genomic Viewer (IGV) screenshots showing H3K27ac marks in the –16 kb region upstream of the *Pparg2* ΤSS in representative wt and *Azin2^-/-^* SAT AP samples. **E,F.** Abundance of H3K27ac (**E**) and H3K9ac (**F**) 251-144 bp upstream of the *Pparg2* ΤSS in differentiated wt and *Azin2^-/-^* SAT APs assessed by ChIP-qPCR and shown as % of input (n=8-10 mice per group). **G.** *Pparg* and *Cepba* expression in differentiated wt and *Azin2^-/-^* SAT APs (n=4-5 mice per group). **H.** Representative images of in vitro differentiated wt and *Azin2^-/-^* SAT APs stained with BODIPY. Scale bar, 200 μm (a representative image of 1 out of 8 mice per group is shown). **I.** Quantification of Oil Red O staining of in vitro differentiated wt and *Azin2^-/-^* SAT APs (n=3 mice per group). **J.** Quantification of Oil Red O staining in SAT APs transfected 3 times during one week of differentiation with siRNA against *Slc7a2* or *Arg1* or non-targeting control siRNA (siCtrl) (n=8 mice per group). **K.** *Pparg* expression in SAT APs transfected 3 times during one week of differentiation with siRNA against *Arg1*, *Sat1*, *Odc1* or non-targeting siRNA (siCtrl) (n=5-7 mice per group). **L.** Quantification of Oil Red O staining in wt and *Azin2^-/-^* SAT APs treated or not with spermidine (SPD, 10 μM) during in vitro differentiation (n=8 mice per group). **M.** Quantification of Oil Red O staining in wt and *Azin2^-/-^* SAT APs treated with C646 (4 μM) or equal amount of DMSO (Ctrl) during in vitro differentiation (n=8 mice per group). **N.** *Pparg* expression in *Azin2^-/-^* SAT APs treated with C646 (4 μM) or DMSO during differentiation (n=5 mice per group). Gene expression in **G, K** and **N** was determined by qPCR using *18S* as a housekeeping gene. All data are shown as mean±SEM. Mann-Whitney U (**E-G,I,K,L,N**), one-way ANOVA (**J,M**) and Kruskal-Wallis (**K** right panel) tests were used for statistical analysis. *p < 0.05, **p < 0.01; ns: not significant

Adipogenesis is initiated with increased expression of C/EBPβ and C/EBPδ, which subsequently cooperate to induce the expression of PPARγ and C/EBPα; the latter activate the transcription of a wide range of factors required for adipocyte maturation ^16^. H3K27ac enrichment of *Pparg* and *Cebpa* gene enhancers indicates their activation and coincides with adipogenic differentiation ^45, 46, 16, 47^. In the *Azin2^-/-^* APs we found enhanced H3K27ac enrichment in the 5’ UTR and intron 2 of the *Pparg2* gene and the *Cebpa* enhancer (distal intergenic region) (*Cebpa* +25.7 kb) (Figure 4D, S4D,F,G). The 5’ UTR of the *Pparg2* gene (starting 60 kb downstream of the *Pparg1* TSS) acts as an enhancer of *Pparg1* during initiation of differentiation and then promotes expression of *Pparg2* encoding for *Pparg* mRNA ^46^. We verified by ChIP-qPCR increased H3K27ac presence in the *Pparg2* promoter in differentiated *Azin2^-/-^*compared to wt APs, using primers binding 251-144 bp upstream of the *Pparg2* TSS (Figure 4E). Moreover, H3K9 acetylation was previously reported to increase in the *Pparg2* promoter during adipogenesis ^48^. Increased H3K9ac enrichment was also detected in the *Pparg2* promoter region (251-144 bp upstream of the *Pparg2* TSS) in differentiated *Azin2^-/-^*compared to wt APs (Figure 4F). Accordingly, *Pparg* and *Cebpa* mRNA expression was elevated in *Azin2^-/-^* SΑΤ APs (Figure 4G). These findings collectively indicate that AZIN2 represses H3K27ac at the enhancers/promoters of key adipogenic and lipid metabolism-associated genes.

We next tested the adipogenic potential of *Azin2^-/-^*APs. Lipid accumulation was significantly increased in in vitro differentiated *Azin2^-/-^* compared to wt SAT APs (Figure 4H,I, S4H,I). *Slc7a2* or *Arg1* siRNA silencing also increased SAT AP adipogenic differentiation (Figure J, S4J, S3G,H). Accordingly, *Arg1*, *Sat1* and *Odc1* mRNA silencing upregulated *Pparg* expression (Figure 4K, S3H-J). These findings indicate that as also shown for polyamine acetylation, AP differentiation depends on arginine uptake and polyamine synthesis. To test the counter idea, that supply of substrates would reduce AP adipogenic differentiation, we added spermidine to the differentiation cultures. Spermidine suppressed AP differentiation in wt but not *Azin2^-/-^* APs (Figure 4L, S4H), standing in accordance with its repressive effect on acetyl-CoA amounts (Figure 3I) and the requirement of AZIN2 for spermidine acetylation (Figure 3A,B,D,E, S3B). H3 acetylation is catalyzed by the lysine acetyltransferase (KAT) p300/CBP (KAT3A/3B) ^47^. Inhibition of p300 by C646 repressed adipogenesis and *Pparg* expression in *Azin2^-/-^* APs (Figure 4M, N). Collectively, these findings suggest that AZIN2-driven polyamine metabolism represses adipogenesis through downregulation of p300-mediated histone acetylation in regulatory elements of key adipogenic genes.

### AZIN2 deficiency leads to increased adipogenesis in vivo

Based on these findings we examined whether AZIN2 deficiency affects adipogenesis in vivo. Chow diet (CD)-fed *Azin2^-/-^* mice had reduced body weight (Figure S5A) but no difference in SAT, GAT, axillary or mesenteric adipose tissue weights compared to littermate wt mice (Figure 5A, S5A-D). Furthermore, there were no differences in liver and spleen weights, glucose tolerance and insulin resistance in *Azin2^-/-^* compared to wt mice (Figure S5E-H). However, SAT and GAT of *Azin2^-/-^* mice contained islets of small adipocytes, which were overall less abundant in wt mice (Figure 5B, S5I). Moreover, the number of total and proliferating (Ki67^+^) APs was lower in SAT of *Azin2^-/-^* compared to wt mice (Figure 5C,D). Based on these data we hypothesize that in young AZIN2 deficient mice shrinkage of the AP pool due to enhanced adipogenesis and limited cell proliferation can predispose to impaired plasticity and dysfunction in the obese state ^4^. In the GAT there was no difference in the number of macrophages (Figure S5J), the percentage of proinflammatory CD11c^+^ macrophages (Figure S5K) and the percentage of profibrotic CD9^high^ APs (Figure S5L) between *Azin2^-/-^* and wt mice.

**Figure 5.**
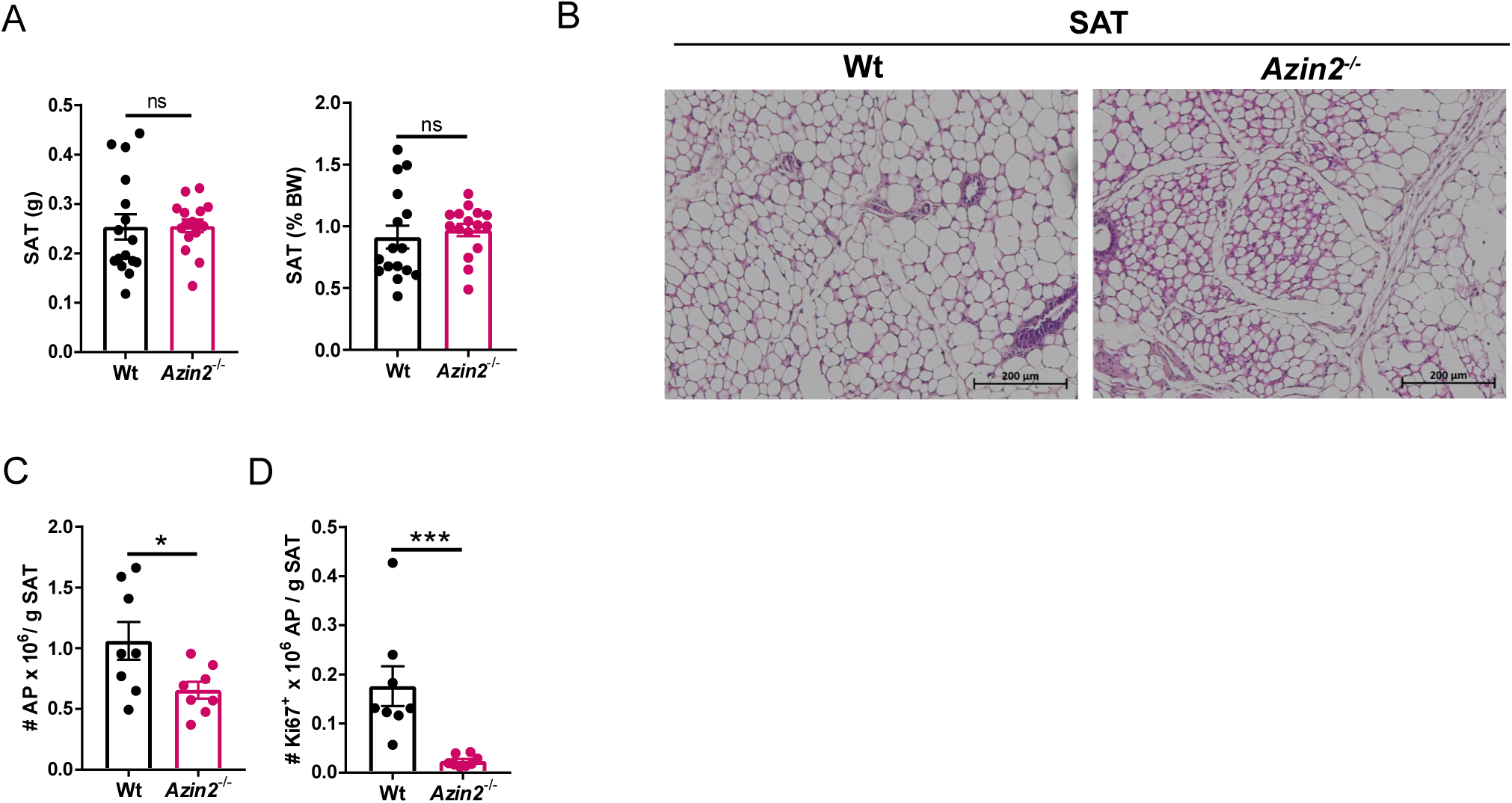
AZIN2 deficiency leads to increased adipogenesis in vivo. **A.** Weight of SAT (left: net weight, right: as percentage of body weight) in 10 weeks-old CD-fed wt and *Azin2^-/-^* mice (n=16 mice per group). **B.** Representative H&E images of SAT in CD-fed wt and *Azin2^-/-^* mice, scale bar: 200 μm (n=3 mice per group). **C,D.** Number of total APs (PDGFRA^+^ LY6A^+^CD45^-^CD31^-^) (**C**) and Ki67^+^ APs (**D**) analyzed by FACS in SAT of 10 weeks-old CD-fed wt and *Azin2^-/-^* mice (n=8 mice per group). Data are shown as mean±SEM. Mann-Whitney U test was used for statistical analysis. *p < 0.05, ***p < 0.001, ns: not significant.

### AZIN2 deficiency leads to increased obesity and adipose tissue dysfunction

We next set out to understand the role of AZIN2 in diet-induced obesity by comparing *Azin2^-/-^* and wt littermate mice fed a HFD (Figure 6A). *Azin2^-/-^*HFD mice gained more body weight and demonstrated reduced insulin sensitivity, energy expenditure and oxygen consumption compared to wt littermate mice (Figure 6B-E) without presenting differences in food intake or locomotion (Figure S6A,B). In accordance, *Azin2^-/-^* HFD mice had increased fat versus lean mass (Figure 6F) and increased SAT mass (Figure 6G) which contained larger adipocytes compared to wt mice (Figure 6H). Notably, the acetyl-CoA concentration in SAT was elevated in *Azin2^-/-^* HFD mice compared to control mice (Figure 6I), consistent with increased acetyl-CoA amounts in *Azin2^-/-^* SAT APs (Figure 3F, S3E). Similar to SAT, GAT was also hypertrophic, consisted of larger adipocytes and contained higher acetyl-CoA amounts in *Azin2^-/-^* compared to wt HFD mice (Figure S6C-F).

**Figure 6.**
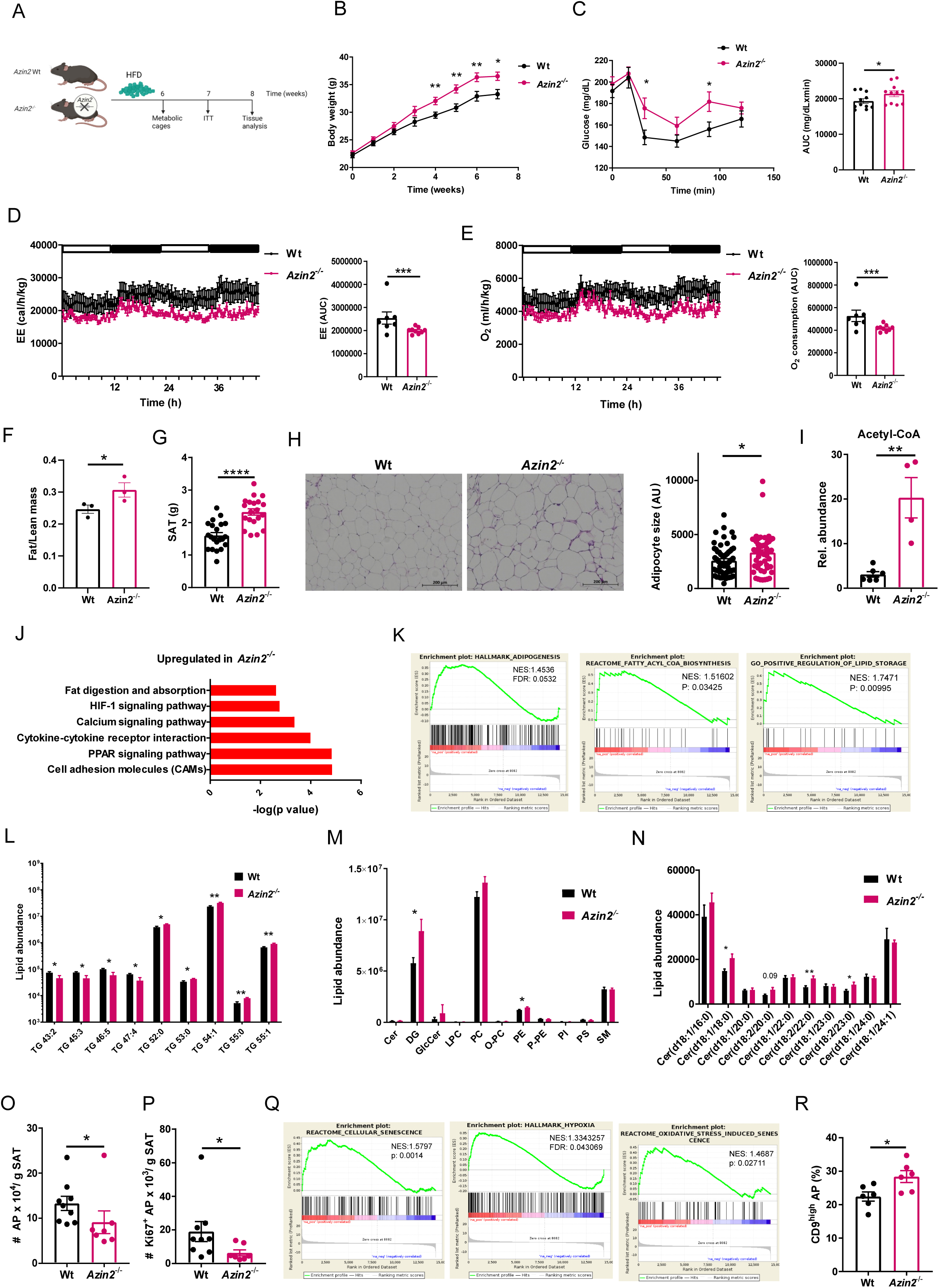
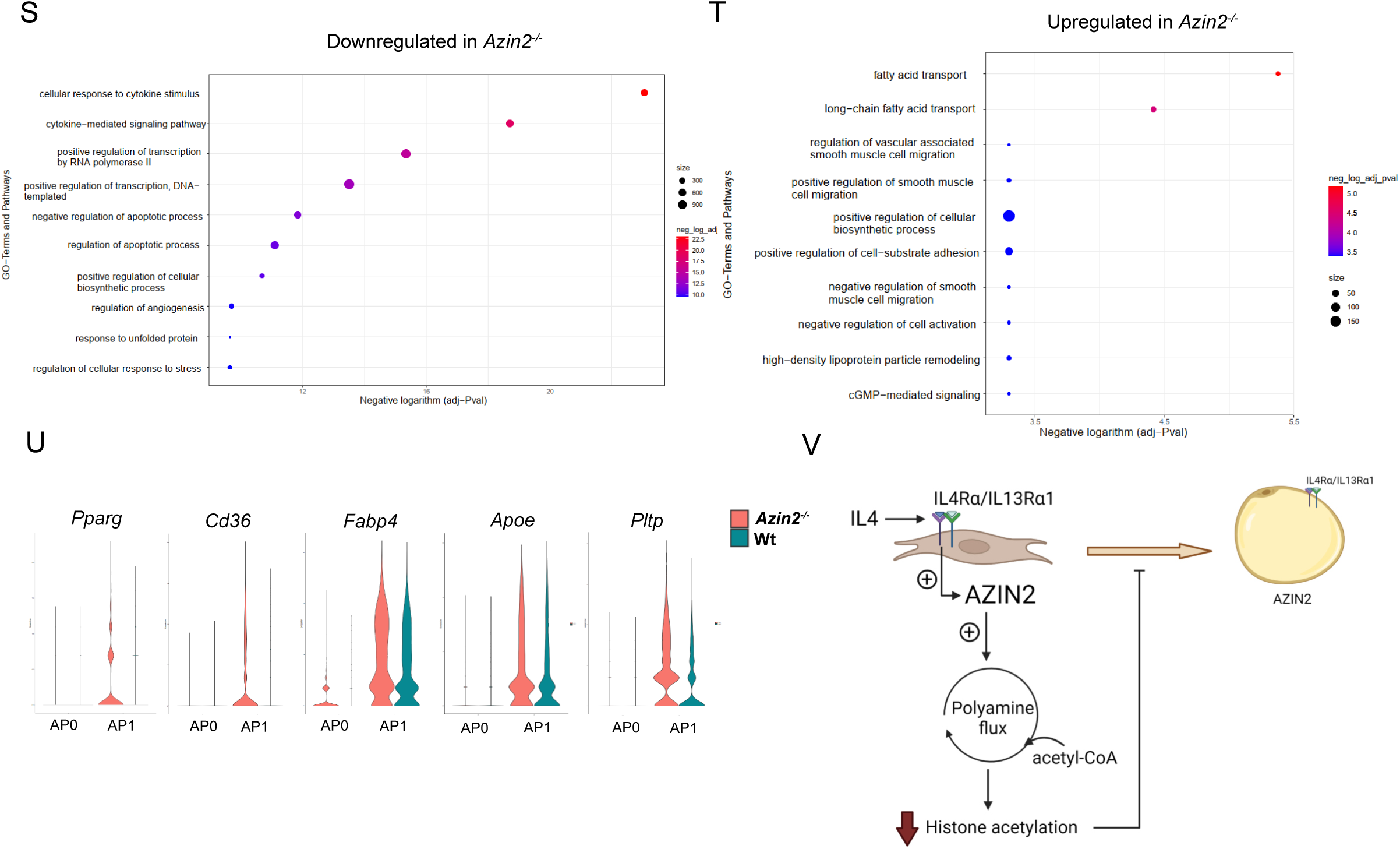
AZIN2 deficiency leads to increased obesity and adipose tissue dysfunction. **A.** Experimental setup. **B.** Body weight gain in wt and *Azin2^-/-^* mice fed for 8 weeks a HFD (n=15 mice per group). **C.** Insulin tolerance test performed at 7 weeks of HFD feeding in wt and *Azin2^-/-^* mice (n=11 mice per group). Left panel: Blood glucose concentration over time; right panel: area under the curve (AUC). **D,E**. Energy expenditure (EE) (**D**) and O_2_ consumption (**E**) measured in metabolic cages in wt and *Azin2^-/-^* mice fed for 6 weeks a HFD (n=7-9 mice per group). **F.** Fat versus lean mass in wt and *Azin2^-/-^*mice fed for 8 weeks a HFD in SAT weight (n=3 mice per group). **G.** SAT weight in wt and *Azin2^-/-^* mice fed for 8 weeks a HFD (n=20-21 mice per group). **H.** Representative images of H&E staining of SAT from wt and *Azin2^-/-^* mice fed for 8 weeks a HFD (one image is shown from one out of 10-11 mice), scale bar: 200 μm. Adipocyte size quantification in SAT of wt and *Azin2^-/-^* mice fed for 8 weeks a HFD (n=6 mice per group, 9-10 sections per mouse were imaged, each dot represents the size of the sliced area of one adipocyte). **I.** Relative abundance of acetyl-CoA levels in SAT of wt and *Azin2^-/-^* mice fed for 8 weeks a HFD presented as peak intensity assessed by LC-MS/MS (n=4-6 mice per group). **J,K.** EGSEA analysis for KEGG signaling pathways upregulated in *Azin2^-/-^* mice (**J**) and GSEA analysis for HALLMARK_Adipogenesis, REACTOME_Fatty Acyl-CoA Biosynthesis and GO_Positive Regulation of Lipid Storage (**K**) based on bulk RNA-seq in SAT of wt and *Azin2^-/-^* mice fed for 8 weeks a HFD (n=4 mice per group). **L-N.** Relative abundance of TG species (**L**), lipid groups (**M**) and ceramide species (**N**) in SAT of wt and *Azin2^-/-^* mice fed for 8 weeks a HFD presented as peak intensity determined by LC-MS/MS, only differentially regulated TG species are shown (n=5-8 mice per group). **O,P.** Total and Ki67^+^ AP (PDGFRα^+^LY6A^+^CD31^-^CD45^-^) abundance per g of tissue in SAT of wt and *Azin2^-/-^* mice fed for 8 weeks a HFD assessed by flow cytometry (n=7-9 mice per group). **Q.** GSEA analysis for REACTOME_Cellular Senescence, HALLMARK_Hypoxia, and REACTOME_Oxidative Stress Induced Senescence pathways based on bulk RNA-seq in SAT of wt and *Azin2^-/-^* mice fed for 8 weeks a HFD (n=4 mice per group). **R.** Percentage of CD9^high^ SAT APs (PDGFRα^+^LY6A^+^CD31^-^CD45^-^) in wt and *Azin2^-/-^* mice fed for 8 weeks a HFD (n=6 mice per group). **S-U.** scRNA-seq was performed in CD45^-^CD31^-^ SAT SVF cells of wt and *Azin2^-/-^* mice fed for 8 weeks a HFD (2 mice pooled per genotype). Down- and up-regulated GO Terms and pathways in *Azin2^-/-^* versus wt mice plotted according to the –log(adjp) value (**S,T**). Violin plots showing gene expression scores of *Pparg*, *Cd36*, *Fabp4*, *Apoe*, and *Pltp*, in AP in wt and *Azin2^-/-^* HFD mice (**U**). Data in **B-F,H,I,L-N,O,P,R** are shown as mean±SEM. Mann-Whitney U test (**B,C,F, I, I,O,P,R**), ANCOVA (**D,E**), t test (**H**) and multiple t test (**L-N**) were used for statistical analysis. *p < 0.05, **p < 0.01, ***p < 0.001, ****p < 0.0001. NES: Normalized enrichment score; FDR: False discovery rate. **V.** Schematic presentation of AZIN2-dependent mechanism regulating adipogenesis.

To further characterize the changes caused by *Azin2* deficiency in fat, we performed bulk RNA-seq in total SAT and GAT of wt and *Azin2^-/-^* HFD mice. In *Azin2^-/-^* HFD mice, pathway analysis showed relative upregulation of pathways linked to fat metabolism, PPAR signaling and inflammatory pathways (Figure 6J) and enrichment of genes related to adipogenesis, fatty-acyl-CoA biosynthesis and lipid storage in SAT (Figure 6K). GAT in *Azin2^-/-^* HFD mice displayed a pronounced inflammatory profile compared to wt counterparts (Figure S6G-L). Expression of many genes involved in inflammation, such as *Tlr8, Fcgr4, Clec12a, C3ar1, Cd68, Fcgr1, Tlr1, S100a8, Igb2, Itgad, Lilrb4a, Trem2, Spp1, Saa3, Itgax, Ccr3, Ccl3* and *Rgs1* was increased in GAT of *Azin2^-/-^* compared to wt HFD mice (Figure S6G). Gene expression related to inflammatory pathways, such as cell adhesion molecules, Toll-like receptor pathway or phagocytosis and *Il6* expression were upregulated and the number of CD11c^+^ ATM and the abundance of F4/80^+^ ATM was increased in GAT of *Azin2^-/-^* HFD mice (Figure S6H-L)^3, 31^. Increased inflammation was accompanied by changes in the metabolic profile of GAT in *Azin2*^-/-^ HFD mice. Gene expression related to oxidative phosphorylation, citrate cycle, pyruvate metabolism and fatty acid degradation were downregulated, while glycolysis and fructose/mannose metabolism were upregulated in GAT in *Azin2^-/-^* HFD mice (Figure S6M,N). In obesity, chronic inflammation in adipose tissue associates with development of fibrosis ^4, 49^. GAT of *Azin2^-/-^* mice showed enhanced expression of a number of fibrotic genes, like matrix metallopeptidase 12 (*Mmp12*), *Mmp13*, collagen, type I, alpha 1 (*Col1a1*), *Col3a1*, *Col8a1* and *Cd9* (Figure S6G,O,P). Accordingly, the abundance of pro-fibrotic CD9^high^ APs was increased in *Azin2^-/-^* HFD mice (Figure S6Q) ^50^. However, we did not observe altered collagen deposition, as the latter was not yet pronounced at 8 weeks of HFD (Figure S6R,S). The above data demonstrate that AZIN2 deficiency leads to adipose tissue inflammation, which is associated with shift to a pro-fibrotic profile, reduced plasticity and impaired lipid storage function of the adipose tissue ^4, 49^.

To analyze the lipidomic profile of the adipose tissue of *Azin2^-/-^* HFD mice, we performed lipidomic analysis ^51, 52^. We found that the total TG acyl chain length was increased and the degree of unsaturation was decreased in SAT of *Azin2^-/-^* mice (Figure 6L, S6T). We previously demonstrated that TG in the adipose tissue get overall longer and more unsaturated with obesity ^51^. Here we show that AZIN2 deficiency impairs this vital adaptation of the adipocyte TG profile to diet-induced obesity. Moreover, total DG was increased in SAT of *Azin2^-/-^* compared to wt HFD mice (Figure 6M). Similarly to TG, DG species upregulated in SAT of *Azin2^-/-^* mice had a low degree of unsaturation, i.e. contained 0-2 double bonds (Figure S6U). DG inhibit insulin signaling and increased DG content in adipose tissue, muscle and liver associates with insulin resistance ^53, 54^. In addition, several ceramide species were more abundant in the SAT of *Azin2^-/-^* compared to wt HFD mice (Figure 6N). The ceramide content in adipose tissue is increased in obesity and type 2 diabetes, and inhibition of ceramide synthesis in adipocytes reduces inflammation and improves insulin sensitivity ^55, 53, 56^. Notably, amounts of C18:0 ceramide, which was reported to increase with obesity in adipose tissue of human subjects ^57, 56^, were increased in SAT of *Azin2^-/-^* HFD mice (Figure 6N). In sum, increased ceramide and DG content and decreased TG desaturation overall demonstrated disturbed lipid storage in the adipose tissue of *Azin2^-/-^* mice.

High AP numbers maintain a healthy adipose tissue by promoting hyperplasia, i.e. formation of many small-sized adipocytes, while low AP numbers associate with adipose tissue hypertrophy, i.e. presence of large adipocytes (Figure 1F, S1K) ^4, 49^. Notably, AP numbers and numbers of proliferating (Ki67^+^) AP were reduced in *Azin2^-/-^*HFD mice (Figure S6O,P), consistent with H3K27ac enrichment in genes linked to negative regulation of cell cycle in *Azin2^-/-^* SAT APs (Figure 4C). Furthermore, genes related to cellular senescence, hypoxia and oxidative stress-induced senescence were enriched in the SAT of *Azin2^-/-^* HFD mice (Figure 6Q). In accordance, senescence assessed by β-galactosidase staining (Figure S7A) and expression of *Cdkn2a* (*p16*) and *Cdkn1a* (*p21*), which encode for negative regulators of cell cycle progression and promote senescence, were increased in *Azin2^-/-^* SAT APs (Figure S7B,C)^62, 63^. In sum, AZIN2 deficiency in obesity leads to increased AP loss, adipocyte hypertrophy, hypoxia and overall adipose tissue dysfunction.

We then investigated the impact of AZIN2 deficiency on the transcriptional profile of APs in SAT of HFD-fed mice by scRNA-seq. CD31^-^CD45^-^ SAT SVF cells of HFD mice consisted of 2 subpopulations, here named AP0 and AP1. Both clusters expressed *Pdgfra* and *Ly6a* (Figure S8A) validating our FACS (PDGFRα^+^LY6A^+^CD31^-^CD45^-^) and bead sorting (CD31^-^CD45^-^) strategies (see above) and standing in accordance with previous reports ^11, 12, 64–66^. Cluster AP0 selectively expressed early AP markers, such as dipeptidylpeptidase 4 (*Dpp4*), wingless-type MMTV integration site family, member 2 (*Wnt2*), bone morphogenetic protein 7 (*Bmp7*) and peptidase inhibitor 16 (*Pi16*) ^11, 12, 14^, dermokine (*Dmkn*) and *Il33* (Figure S8B), while the AP1 cluster exhibited higher expression of adipogenic genes, such as lipoprotein lipase (*Lpl*) and fatty acid binding protein 4 (*Fabp4*), coagulation factor III (*F3*/*Cd142*) and genes encoding proteins linked to ECM remodeling, such as biglycan (*Bgn*), lumican (*Lum*), matrix Gla protein (*Mgp*), *Mmp3* and transforming growth factor, beta 1 (*Tgfb1*) compared to AP0 cells (Figure S8C and 6U). These data indicated that the AP0 cluster consisted of early progenitors, while AP1 progenitors expressed markers of adipogenesis. This clustering stands in accordance with previous studies ^11, 12, 14, 66, 15^: AP0 corresponds to P1 of Schwalie et al. ^15^, ASC2 of Burl et al. ^11^, group 1 of Merrick et al. ^14^, FAP3 of Sarvari et al., ^66^ and mASPC2 of Emont et al., ^12^, and AP1 corresponds to P2 of Schwalie et al. ^15^, ASC1 and differentiated ASC of Burl et al. ^11^, group 2 of Merrick et al. ^14^, FAP2 of Sarvari et al., ^66^ and mASPC1 of Emont et al., ^12^. These two clusters also correspond to previously reported human AP subpopulations ^12, 14^.

*Azin2* was expressed in both AP0 and AP1 (Figure S8D). Pathway analysis showed that gene expression related to cellular response to cytokines, gene transcription and negative regulation of apoptosis were downregulated, while gene expression related to lipid metabolism and lipid transport was upregulated in APs of *Azin2^-/-^* compared to wt mice (Figure 6S,T). Specifically, expression of *Pparg*, *Cd36, Fabp4*, apolipoprotein e (*Apoe*) and phospholipid transfer protein (*Pltp*) was upregulated in AP1 of *Azin2^-/-^* mice (Figure 6U). In contrast, expression of genes encoding for anti-oxidant proteins, such as metallothionein 2 (*Mt2*), proteins mediating interactions with the ECM, such as hyaluronan synthase 1 (*Has1*), UDP-glucose 6-dehydrogenase (*Ugdh*), *Cd44* and pentraxin 3 (*Ptx3*), and proteins mediating the response to mitogenic signals, such as immediate early response 5 (*Ier5*) and lamin a (*Lmna*), was downregulated in both clusters in *Azin2^-/-^* compared to wt mice (Figure S8E). Notably, expression of *Il4ra*, which was detected only in the AP1 cluster and found to be a negative regulator of adipogenesis and inducer of *Azin2* expression (Figure 2B,D,F-I, S2C,F)^38, 39^, was decreased in *Azin2^-/-^* mice (Figure S8E). Collectively, these data suggest that AZIN2 modulates AP identity, while its deficiency leads to increased pre-adipogenic commitment and reduced interaction with the microenvironment, decreased response to mitogenic signals, and reduced production of antioxidant agents. Mechanistically, IL4 promotes AZIN2 expression in APs, which regulates acetyl-CoA levels and adipogenesis-related gene expression through histone acetylation (Figure 6V).

### AZIN2 deficiency leads to enhanced liver steatosis in HFD

Prolonged obesity leads to steatohepatitis ^67^. We asked whether AZIN2 deficiency affects liver steatosis in HFD mice. Strikingly, *Azin2^-/-^* mice fed for 8 weeks a HFD had increased liver weight (Figure S9A), enhanced steatosis and triglyceride content (Figure S9B,C), and increased expression of lipogenesis-related genes, such as *Pparg*, *Cebpa*, *Cd36*, *Acc* and *Fasn*, and fibrosis-related genes, such as *Col1a1* and *Col3a1*, compared to wt control mice (Figure S9D). Expression of inflammatory genes, such as *Il6*, *Tnf* or *Tgfb*, was not altered (not shown). Given the low *Azin2* expression in the liver and hepatocytes (Figure S1I,J), we hypothesize that the effects of the AZIN2 deficiency in the liver derive from its effects on the adipose tissue (Figure 6).

### *AZIN2* expression negatively correlates with adipogenesis markers in the human adipose tissue

Finally, we mined the GEPIA2 database ^41^ for expression of genes encoding for early progenitor, adipogenesis and mature adipocyte markers in correlation with *AZIN2* expression in human adipose tissue. *AZIN2* mRNA expression positively correlated with expression of the early progenitor markers *DPP4* and *WNT2* (Figure S10A,B,R,S), *PDGFRA* (Figure S10C) and *EPHA3*, a marker of anti-adipogenic progenitors (Figure S10D,T), while it negatively correlated with expression of genes encoding for commitment markers, like *F3*/*CD142* (Figure S10E,U), adipogenesis markers, such as *FABP4*, *PPARG* and *ADIPOQ* (Figure S10F-H,V-X), adipocyte markers, like patatin-like phospholipase domain containing 3 (*PNPLA3*), diacylglycerol O-acyltransferase 2 (*DGAT2*), sterol regulatory element binding transcription factor 1 (*SREBF1*) and post-GPI attachment to proteins inositol deacylase 1 (*PGAP1*) (Figure S10I-L, Y), and genes involved in lipid metabolism, including ELOVL fatty acid elongase 5 (*ELOVL5*), *ELOVL3*, and fatty acid desaturase 3 (*FADS3*) (Figure S10M,N,Z,AA) ^12, 14, 66, 15^. *AZIN2* expression also positively correlated with the expression of endothelin receptor type A (*EDNRA*), lin-7 homolog A, crumbs cell polarity complex component (*LIN7A*), alkylglycerol monooxygenase (*AGMO*) and formin homology 2 domain containing 3 (*FHOD3*), which were previously identified as markers of an adipocyte subpopulation with protective function against insulin resistance (Figure S10O-Q,AC,AD)^12^. These correlations collectively suggest that as in the mouse, in humans *AZIN2* expression associates with features of healthy adipose tissue.

## Discussion

Maintenance of adipose tissue homeostasis requires constant adipocyte renewal through AP differentiation ^68^. In periods of increased energy intake or reduced energy expenditure lipid storage in the adipose tissue protects against ectopic lipid deposition and lipotoxicity; however, excessive adipose tissue expansion is associated with increased adipocyte destruction, hypoxia, inflammation, fibrosis and overall loss of adipose tissue plasticity, which in turn promotes dyslipidemia, systemic inflammation and insulin resistance ^3, 4^. In addition, APs can combust energy through thermogenesis, especially in response to cold or exercise, and thus have a protective function against obesity and associated dysmetabolism. At the beginning of obesity, APs differentiate to adipocytes, which then grow in size by storing TGs ^6, 7^. Hence, formation of new adipocytes through adipogenesis is required for adipose tissue expansion. Moreover, we suggest that excessive adipogenesis in the lean state or young age may cause AP depletion leading to aggravated adipose tissue pathophysiology in obesity. This does not contradict the fact that in established obesity adipose tissue hyperplasia favors metabolic advantage compared to adipose tissue hypertrophy ^4^.

While the molecular mechanisms regulating gene transcription during adipogenic differentiation have been dissected in detail ^69, 70, 16^, little is known about the adipose tissue microenvironment-derived signals regulating AP differentiation. Moreover, how cell metabolic circuits regulate adipogenesis was so far unclear. Here, we show that polyamine metabolism is downregulated in the adipose tissue upon diet-induced obesity. Components of polyamine metabolism including AZIN2 are expressed in APs and are downregulated upon adipogenic differentiation. Moreover, IL4 upregulated AZIN2 expression, in keeping with its anti-obesogenic effect ^33, 38, 39^. Accordingly, in the human adipose tissue *AZIN2* expression positively correlates with expression of type 2 immunity cytokines.

We found that AZIN2 promotes polyamine synthesis and acetylation in APs. Similarly, AZIN2 was recently linked to increased amounts of acetylated polyamines in the brain in tau neuropathy ^71^. In accordance, AZIN2 negatively regulates acetyl-CoA amounts in APs and the adipose tissue. Elevated acetyl-CoA amounts in *Azin2^-/-^* APs may drive increased H3K27ac in adipogenic genes, including *Pparg* and *Cebpa*, which encode for transcription factors with pivotal roles in adipogenic commitment ^16^. AZIN2 deficiency leads to enhanced adipogenesis and AP depletion, while it also associates with AP senescence, thereby predisposing for development of an unhealthy adipose tissue upon obesity. On the other hand, increased acetyl-CoA amounts in the adipose tissue of *Azin2^-/-^* mice are linked to altered TG desaturation, and increased ceramide and DG content, i.e. perturbed lipid storage, which collectively can drive lipotoxicity, inflammation and insulin resistance ^55, 53, 54, 56, 49, 72^. Indeed, *Azin2^-/-^* HFD mice exhibit increased adipose tissue inflammation and systemic insulin resistance compared to wt HFD mice.

At the single-cell level, AZIN2 deficiency increases expression of genes related to adipogenesis and lipid metabolism and reduces expression of genes encoding for factors mediating cell-to-matrix interactions in committed (AP1) progenitors, inferring that adipogenic differentiation associates with reduced interaction of APs with their microenvironment. *Il4ra* expression is diminished in *Azin2^-/-^* AP1 cells, standing in accordance with its reduction upon adipogenesis, and suggesting that *Azin2^-/-^* AP1 cells are less responsive to the anti-adipogenic effects of IL4, which in turn may further accelerate their differentiation to adipocytes ^73, 38, 39^. Similarly, in human adipose tissue, *AZIN2* expression positively correlates with early progenitor markers and negatively correlates with expression of adipogenesis and lipid processing markers ^12, 14, 66^ To which extent AZIN2 plays a protective role in human obesity will be examined in future studies. Furthermore, AZIN2 was found to maintain a ‘healthy’ AP state as its deficiency predisposes to reduced AP numbers and AP senescence. In support of this, genes related to senescence and cell cycle regulation present increased H3K27ac marks in *Azin2^-/-^* APs, suggesting inhibition of senescence by AZIN2 through epigenetic regulation. Previous reports showed that polyamine depletion leads to hyperacetylation, ROS generation and necrosis, while spermidine treatment induces H3 deacetylation through inhibition of histone acetyltransferases and suppresses oxidative stress in yeast ^20^. Accordingly, we show that spermidine decreases acetyl-CoA amounts in APs by fueling SAT1, which catalyzes spermidine acetylation. SAT1 was previously shown to be a regulator of acetyl-CoA levels in adipose tissue and SAT1 overexpression or knockout reduced or increased body fat in HFD fed mice, respectively ^74–78^. Our work indicates that AZIN2 determines SAT1 activity in APs.

Spermidine and spermine were previously shown to protect against diet-induced obesity and associated metabolic disturbances ^79–84^. We show that spermidine inhibits adipogenesis in wt but not AZIN2 deficient APs indicating that spermidine acetylation and not spermidine per se inhibits adipogenesis. Hence, spermidine supplementation is not expected to revert increased obesity in AZIN2 deficient mice, as it does in wt mice ^79–84^. Spermidine also promotes autophagy, mitophagy and mitochondrial respiration in cardiomyocytes, thereby providing cardioprotection ^19^, while it inhibits oxidative stress and extends the lifespan of cells and organisms spanning from yeast and worms to mice and humans ^19, 20, 85^. Recently, spermidine was shown to activate fatty acid oxidation in CD8^+^ T cells through direct binding to and activation of mitochondrial trifunctional protein (MTP) ^86^. Moreover, spermidine is a precursor of hypusine, a post-translational modification of eukaryotic initiation factor 5A isoform 1 (eIF5A), which promotes the expression of proteins involved in mitochondrial metabolism in macrophages ^60^. In accordance with these effects of spermidine, AZIN2 preserves mitochondrial metabolism, i.e. TCA cycle function, oxidative phosphorylation and lipid oxidation, and reduces oxidative stress in the adipose tissue of obese mice. Whether AZIN2 also counteracts aging and extends the lifespan of cells and organisms merits further investigation.

The characteristics of different adipose tissue depots can vary significantly depending on the anatomical position. In humans, APs are more abundant in gluteofemoral compared to abdominal subcutaneous and omental adipose tissue ^87^. This is in agreement with the rather hyperplastic nature of gluteofemoral subcutaneous compared to visceral adipose tissue and the metabolically favorable fat deposition at the gluteofemoral area compared to the visceral site ^4^. On the other hand, adipose tissue hyperplasia in gonadal and inguinal subcutaneous adipose tissue differs in males and females ^9, 6, 7^. Sex-dependent adaptations of polyamine metabolism or other cell metabolic circuits in different adipose tissue depots is an important open question, which merits future investigation.

In conclusion, this study demonstrates that AZIN2 regulates acetyl-CoA amounts, histone acetylation and transcriptional networks that determine AP viability, identity and function (Figure 5AA). Inactivation of AZIN2 exacerbates adipose tissue growth, associated with adipocyte hypertrophy, and enhances adipose tissue inflammation and fibrosis, and systemic insulin resistance in obesity. Our findings have important implications for understanding and potentially therapeutically interfering with obesity development.

## Acknowledgments

We thank Marta Prucnal for the technical assistance. We also thank the Core Facility Cellular Imaging of the Medical Faculty Carl Gustav Carus, TU Dresden, especially Michael Gerlach, for the technical support. This work was supported by funds from the Deutsche Forschungsgemeinschaft (AL 1686/6-1 and SFB-TRR 205 project A07 to VIA and SFB-TRR 127 to TC), the ‘Sonderzuweisung zur Unterstützung profilbestimmender Struktureinheiten 2023-2024’ by the SMWK (Saxon State Ministry of Science, Culture and Tourism) to ÜC and TC and the ‘Sonderzuweisung zur Unterstützung profilbestimmender Struktureinheiten 2021’ by the SMWK and TG70 by Sächsische Aufbaubank and SMWK (the measure is cofinanced with tax funds on the basis of the budget passed by the Saxon State Parliament) to MF.

## Author contributions

**C.M.:** Investigation; **A.S.:** Software, formal analysis, data curation; **A.A.**: Methodology, validation, investigation; **I.M.:** Methodology, validation, investigation; **E.H.:** Investigation; **S.T.:** Methodology, investigation, data curation; **B.G.:** Investigation; **A.S.:** Investigation; **J.P.:** Investigation; **L.A.:** Investigation; **M.W.:** Investigation, formal analysis, data curation; **N.W.:** Investigation; **G.F.:** Investigation; **P.S.:** Investigation; **KJ.C:** Methodology; **S.G.:** Investigation; **M.L.:** Investigation; **M.E.:** Resources; **T.G.:** Resources; **B.W.:** Resources; **A.D.:** Software, formal analysis; **M.G.:** Resources; **M.B.:** Editing; **Ü.C.:** Resources; **D.V.:** Resources; **M.F.:** Resources; **M.P.:** Resources, formal analysis, data curation; **P.M.:** Methodology, editing; **T.C.:** Conceptualization, resources, editing, funding acquisition. **V.I.A.:** Conceptualization, validation, resources, data curation, writing-original draft, editing, supervision, project administration, funding acquisition.

## Data and code availability

The bulk RNA-seq, ChIP-seq and scRNA-seq data reported in this paper are accessible in the reference Series GSE235012 under the link https://www.ncbi.nlm.nih.gov/geo/query/acc.cgi?acc=GSE235012. This paper does not report original code. Any additional information required to reanalyze the data reported in this paper is available from the lead contact upon request.

## STAR Methods

### Mouse lines

For the generation of *Azin2^-/-^* mice, zygotes were isolated from superovulated female C57Bl/6NCrl mice at E0.5. Single zygotes were microinjected with Cas9 RNPs prepared from 333.3 ng/μl NLS-Cas9 (ToolGen), 1 μM gRNA1 (5’TCAGCGACGCCCCAAACCTC3’), 1 μM gRNA2 (5’GAGGGCGGCGGCACATTCAC3’), and 2 μM tracrRNA (Integrated DNA Technologies). Embryos that developed to the two-cell stage were transferred into pseudopregnant foster mothers. *Azin2^-/-^* mice carried deletion of *Azin2* exons 6 to 9. PCR genotyping was used to identify founders with mutated alleles. PCR genotyping was performed using following primers for the *Azin2^-/-^* Fw 5’ACCTGCCTGGTATGGAGTTG3’, Rv 5’GTGCCTCTGCCTCCTAAGTG3’ (product band size 575 bp) and wt mice: Fw 5’GCTCTTCCTCCTTTCCCAGT3’, Rv 5’CCACGCTTGACCACACATTA3’ (product band size 206 bp).

### In vivo experiments

All feeding experiments were performed in male mice and were started at the age of 6-7 weeks. C57BL/6J wt male mice were fed for 20 weeks a LFD (10% of kcal from fat, D12450B, Research Diets) or a HFD (60% of kcal from fat, D12492, Research Diets). *Azin2^-/-^* and wt littermates were fed for 8 weeks a LFD or a HFD. The body weight was measured every week during the course of the feeding. For glucose tolerance test (GTT) mice were fasted overnight and then were intraperitoneally (i.p.) injected with -(+)-glucose (Sigma Aldrich) (1 g/kg of body weight). For insulin tolerance tests (ITT), mice were fasted for 6 h and then i.p. injected with 0.5 or 1 U/kg insulin (Eli Lilly and Company) in lean and obese mice, respectively, and blood glucose levels were measured via the tail vein with an Accu-Chek glucosemeter (Roche) at 0, 15, 30, 60, 90 and 120 min ^29, 30^. The metabolic profile of mice was analyzed in metabolic cages as previously described ^29^. Briefly, mice of similar weights were individually housed in metabolic cages (PhenoMaster, TSE Systems) for 3 days with free access to water and food maintaining a 12 h:12 h light/dark cycle. A period of at least 24 h acclimatization in the metabolic cages preceded initiation of the experiment and data collection. Volume of oxygen consumption (VO_2_), carbon dioxide production (VCO_2_), food intake and locomotion were determined every 20 minutes. Energy expenditure (EE) was calculated as 3.941xVO_2_+1.106xVCO_2_.

For IL4 treatments mice were fed for 6 weeks a HFD and during the last 2 weeks of HFD feeding they were i.p. injected every second day with a complex of mouse recombinant IL4 (2 μg/mouse) (Peprotech) and anti-IL4 (10 μg/mouse) (Tonbo Biosciences) diluted in PBS as previously described ^88, 32^. Control mice received equal volume of PBS i.p. injections.

Mice were injected with 500 L3-stage *N. brasiliensis* larvae subcutaneously and analyzed 9 days later, as previously described ^89^.

At the end of the experiments, inguinal SAT and GAT were collected and either snap-frozen or further processed for SVF or AF isolation or cell sorting. Animal experiments were approved by the Landesdirektion Sachsen, Germany, the Regierung von Mittelfranken, Germany and the Bezirksregierung Leipzig, Bezirksregierung Halle, Germany.

### AP isolation

SAT and GAT were harvested, minced on ice and incubated for 45 min with collagenase type I (Thermo Fisher Scientific) (2 mg/ml/g tissue) at 37 °C under continuous rotation. Cells were then passed through a 100-μm pore-sized cell strainer and centrifuged at 300 g for 10 min. In some experiments the floating AF was collected. Erythrocytes were removed from the SVF through lysis with a red blood cell (RBC) lysis buffer (Biolegend) and SVF cells were used for RNA isolation, FACS analysis or AP sorting. APs were sorted by FACS as PDGFRα^+^LY6A^+^CD31^-^CD45^-^ cells with a BD FACS ARIA apparatus (BD Biosciences) and using the BD FACSDIVA v8.0.1 software (BD Biosciences) ^29^ or as CD31^-^CD45^-^ cells using anti-CD31 and anti-CD45 MicroBeads (Miltenyi Biotec), according to manufacturer’s instructions. CD31^+^ and CD45^+^ cells were also kept and analysed. In some experiments CD11b^-^ cells were isolated from SAT SVF with anti-CD11b MicroBeads (Miltenyi Biotec).

### Cell and explant culture

APs were grown for one week in 4.5 g/L glucose Glutamax–DMEM with 10% FBS and 1% penicillin/streptomycin at 37 °C and 5% CO_2_. For adipogenic differentiation they were cultured in medium containing 5 µg/ml insulin (Eli Lilly and Company), 0.5 mM 3-isobutyl-1-methylxantine (Sigma-Aldrich), 1 µM dexamethasone (Sigma-Aldrich) and 1 µM rosiglitazone (Sigma-Aldrich). After 2 days, they were switched to medium containing 5 µg/ml insulin and cultured for another 5 days. Medium was changed every 2 days. To study senescence, APs were differentiated for one week and cultured for another week in culture medium.

3T3-L1 pre-adipocytes (CL-173, ATCC) were maintained in 4.5 g/L glucose Glutamax – supplemented DMEM with 1% FBS and 1% penicillin/streptomycin at 37 °C and 5% CO_2_.

Human pre-adipocytes (P10761, Innoprot) were cultured for up to 3 passages in preadipocyte medium containing 1% preadipocyte growth supplement, 5% FBS and 1% penicillin/streptomycin (Innoprot) at 37 °C and 5% CO_2_ on cell culture plates pre-coated with 2 μg/cm^2^ poly-L-lysine. For differentiation, they were cultured for 2 days in differentiation medium containing 1% preadipocyte differentiation supplement, 5% FBS and 1% penicillin/streptomycin (Innoprot) and for another 6 days in preadipocyte medium containing 0.5 mg/ml insulin.

SAT or GAT explants from *Azin2^-/-^*, *Il4ra^-/-^* and their respective wt littermate control mice were isolated, minced and kept for 18 h in 5 g/L glucose Glutamax – supplemented DMEM with 1% FBS and 1% penicillin/streptomycin at 37 °C and 5% CO_2_.

### Cell and explant treatments

Undifferentiated APs were treated for 4 or 24 h with mouse recombinant IL4 or IL13 (both at 20 ng/ml, purchased from PeproTech) in 1% FBS 4.5 g/L glucose Glutamax – supplemented DMEM. Undifferentiated human preadipocytes were treated for 24 h with 12 ng/ml human recombinant IL4 (PeproTech). During differentiation APs were treated with 10 μM spermidine (Sigma-Aldrich), 4 μM C646 (Sigma-Aldrich) or respective controls. All agents were re-added every 2 days with fresh medium. For arginine metabolite measurement, cells were starved for 2 h in Silac medium (Thermo Fisher Scientific) without FBS containing 146 mg/l L-lysine (Sigma-Aldrich), then 400 μΜ arginine (Sigma-Aldrich) was added and cell lysates or supernatants were collected after 60 min.

### siRNA transfections

Undifferentiated APs were transfected with TARGETplus SMARTpool siRNA against *Azin2*, *Slc7a2*, *Arg1*, *Sat1*, *Odc1*, *Stat6* or control non-targeting siRNA (all at 30 nM and from Horizon Discovery) using Lipofectamine RNAiMAX transfection reagent (Thermo Fisher Scientific) and the reverse transfection method per manufacturer’s instructions. During differentiation, siRNA transfections were repeated every 2-3 days using same siRNA concentration and Lipofectamine volume.

### Plasmid transfection

3T3-L1 cells were transfected with a plasmid overexpressing *Azin2* (NM_172875, mouse tagged ORF clone, Origene) or pCMV control plasmid using Lipofectamine LTX Reagent (Thermo Fisher Scientific) according to manufacturer‘s instructions and as previously described ^90^.

### Bulk RNA-seq

Bulk RNA-seq was performed and analyzed as previously described ^91, 52^. For transcriptome mapping, strand-specific paired-end sequencing libraries from total RNA were constructed using TruSeq stranded Total RNA kit (Illumina Inc). Sequencing was performed on an Illumina HiSeq3000 (1×75 basepairs). Low quality nucleotides were removed with the Illumina fastq filter and reads were further subjected to adaptor trimming using cutadapt ^92^. Alignment of the reads to the mouse genome was done using STAR Aligner ^93^ using the parameters: “–runMode alignReads –outSAMstrandField intronMotif –outSAMtype BAM SortedByCoordinate -- readFilesCommand zcat”. Mouse Genome version GRCm38 (release M12 GENCODE) was used for the alignment. The parameters: ‘htseq-count -f bam -s reverse -m union -a 20’, HTSeq-0.6.1p1 ^94^ were used to count the reads that map to the genes in the aligned sample files. The GTF file (gencode.vM12.annotation.gtf) used for read quantification was downloaded from Gencode (https://www.gencodegenes.org/mouse/release_M12.html). Gene centric differential expression analysis was performed using DESeq2_1.8.1 ^95^. The raw read counts for the genes across the samples were normalized using ‘rlog’ command of DESeq2 and subsequently these values were used to render a PCA plot using ggplot2_1.0.1 ^96^.

Pathway and functional analyses were performed using GSEA ^96^ and EGSEA ^97^. GSEA is a stand-alone software with a graphical user interface (GUI). To run GSEA, a ranked list of all the genes from DESeq2 based calculations was created using the -log10 of the p-value. This ranked list was then queried against Molecular Signatures Database (MsigDB), KEGG, GO, Reactome and Hallmark based repositories. EGSEA is an R/Bioconductor based command-line package. For pathway analysis with EGSEA lists of DEG with padj < 0.05 or p < 0.05 were used based on the KEGG database repository. For constructing heatmaps, the “rlog-normalized” expression values of the significantly expressed genes (padj < 0.05) was scaled using z-transformation. The resulting matrices were visually rendered using MORPHEUS ^91, 52^.

### scRNA-seq

CD45^-^CD31^-^ SVF cells were isolated by negative selection with MicroBeads (Miltenyi Biotec) from SAT of *Azin2^-/-^*and wt mice (2 mice pooled per genotype) fed for 8 weeks a HFD. 10,275 wt and 11,004 *Azin2^-/-^* cells were subjected to droplet-based scRNA-seq. Specifically, the sorted cells were concentrated by centrifugation (300 rcf, 5 min, 4 °C) and the volume was reduced to 42 µl. Cells were carefully resuspended, cell concentration and quality was determined and cells were added to the reverse transcription mix before loading cells on the 10X Genomics Chromium system in a Chromium Single Cell G Chip and processed further following the guidelines of the 10x Genomics user manual (v3.1). In short, the droplets were directly subjected to reverse transcription, the emulsion was broken and cDNA was purified using Silane beads. After the amplification of cDNA with 11 cycles, it underwent 0.6x SPRI bead purification and quantification. The 10X Genomics scRNA-seq library preparation – involving fragmentation, dA-Tailing, adapter ligation and a 12 cycles indexing PCR – was performed with 25% of the cDNA material based on the manufacturer’s protocol. After quantification, the libraries were sequenced on an Illumina Novaseq6000 in paired-end mode (R1/R2: 100 cycles; I1/I2: 10 cycles), thus generating ∼300 million fragment pairs. The raw sequencing data was then processed with the ‘count’ command of the Cell Ranger software (v6.1.2) provided by 10X Genomics with the option ‘--expect-cells’ set to 10,000 (all other options were used as per default). To build the reference for Cell Ranger, mouse genome (GRCm39) as well as gene annotation (Ensembl 104) were downloaded from Ensembl. Genome and annotation were processed following the build steps provided by 10x (https://support.10xgenomics.com/single-cell-gene-expression/software/release-notes/build#mm10_2020A) to build the appropriate Cellranger reference.

Seurat (v 4.3.0)^98^ was used to perform analyses of the scRNA-seq data in R (v 4.0.2) environment. Briefly, wt sample (10,275 cells and 17,032 genes) and *Azin2^-/-^* sample (11,004 cells and 17,476 genes) were analyzed using the SCTransform command (running on parameters: vst.flavor = "v2", verbose = FALSE, ncells=10,000, variable.features.n = 6,000). Subsequently, RunPCA, RunUMAP and FindNeighbors commands were run using 50 dimensions. A “resolution” of 0.07 was used to run FindClusters command. The resultant Seurat “objects” were integrated as described on https://satijalab.org/seurat/articles/sctransform_v2_vignette.html. The numbers of features and dimensions used were same as above. The “resolution” was set to 0.03. NormalizeData was used to perform normalization on the integrated dataset with normalization.method set to "LogNormalize" and scale.factor set to 10,000. FindVariableFeatures, with “selection.method” set to "vst" and number of features limited to 6,000 was used to find the most variable genes. ScaleData command was then used to run the Seurat object using the variable features found in the previous step. Subsequently, “FindMarkers” command was used to find the differentially expressed genes, with logfc.threshold set to 0.1., between the *Azin2^-/-^* and wt cells. The R package enrichR (which provides an API to https://maayanlab.cloud/Enrichr/ webserver ^99^) was used to perform functional analyses of the differentially expressed genes. Plotting was done using the functions from Seurat and ggplot2.

### ChIP-qPCR

ChIP was performed using the EZ-Magna ChIP HiSens (Millipore) following manufacturer’s instructions. Briefly, differentiated APs were cross-linked using 1% formaldehyde for 10 min at room temperature (RT) and quenched with glycine. Chromatin shearing was achieved by sonication for 25 cycles (20 sec on and 20 sec off) on wet ice to generate DNA fragments with average length of 200-500 bp. Five µl of sheared cross-linked chromatin were kept to be used as 100% input. Immunoprecipitation was performed by incubating the sheared chromatin with the magna ChIP protein A/G magnetic beads and the antibody either against H3K27Ac (Millipore 17-683), H3K9Ac (Millipore 17-658) or IgG control antibody overnight at 4 °C. Histone acetylation of *Pparg2* promoter was assessed by qPCR using the following primers: F 5’TGTGTGGGTCACTGGCGAGACA3’; R 5’TGGCTGGCACTGTCCTGACTGA3’ and the SsoFast Eva Green Supermix (Bio-Rad, Munich, Germany), a CFX384 real-time System C1000 Thermal Cycler (Bio-Rad) and the Bio-Rad CFX Manager 3.1 software.

### Bulk ChIP-seq

For ChIP-seq, libraries were generated using the NEBNext Ultra II DNA library Prep kit (Illumina Inc). Nextflow (v 22.02.1) ^100^ was used to run the nf-core/chipseq: 2.0.0 workflow pipeline (https://nf-co.re/chipseq) with the following parameters : --genome GRCm38 --macs_pvalue 0.1 --max_cpus 4 --macs_gsize 1.87e9.The “narrowPeak” files generated from the pipeline were further numerically sorted by the region coordinates and then merged (using the “merge” function from bedtools v2.17.0 ^101^) to create a consensus bed file. This file was subsequently used along with “featureCounts” ^102^(running on parameters: isPairedEnd=TRUE, checkFragLength=TRUE, requireBothEndsMapped=TRUE) to create counts matrix for all the samples. Normalization and differential analyses were performed using DESeq2 ^95^. Functional annotation of the differentially activated regions was done using g:Profiler webserver ^103^. Figures were plotted using ggplot2 in R environment ^96^. “bigwig” files were used to visualize genomic regions in IGV ^104^. Following commands from Deeptools package ^105^ executed sequentially using the listed respective parameters to generate the profile and heatmap plots for the ChIP-Seq data: 1) bamCompare --binSize 20 --normalizeUsing BPM --smoothLength 60 --extendReads 150 --centerReads -- scaleFactorsMethod None 2) computeMatrix scale-regions --beforeRegionStartLength 5,000 -- regionBodyLength 5,000 --afterRegionStartLength 5,000 --skipZeros (a bed file listing the differentially activated regions were provided) 3) plotProfile –perGroup red red red blue blue blue -out --samplesLabel "KO1" "KO2" "KO3" "WT1" "WT2" "WT3" --refPointLabel "TSS" -- yAxisLabel "Average Signal Intensity" 4) plotHeatmap2 -samplesLabel "KO1" "KO2" "KO3" "WT1" "WT2" "WT3"--regionsLabel "Genome-Wide Assay Signal at Genes" --yAxisLabel "Fold Enrichment (Average Log2 IP/Input)" --zMax 0.5 --legendLocation none.

### Arginine metabolite measurement

Arginine and its metabolites ornithine, putrescine, spermidine, spermine, N1-acetylspermidine, and N1-acetylspermine were measured by liquid chromatography-tandem mass spectrometry (LC-MS/MS) as previously described ^106^. In brief, cultured cells were washed with cold PBS and lysed in the culture plates using methanol/water (1/1) (200 μl per well in 24-well plates). Adipose tissue was homogenized in 200 μl acetonitrile/water (1/1) using a tissue grinder. Ten µl of cell culture supernatant were mixed with 200 µl acetonitrile/water (1/1). Samples were vortex-mixed for one minute and centrifuged at 3,000 g for 10 min at 4 °C. Supernatants from extracted samples were transferred directly onto a 96-well polytetrafluoroethylene (PTFE)-filter plate (Merck-Millipore) and filtered by assistance of positive pressure. Subsequently, filtered extracts were dried in a vacuum-assisted centrifuge, thereafter reconstituted in 200 µl initial mobile phase and analyzed by LC-MS/MS.

### Lipidomics

Lipids from approximately 50 mg adipose tissue were extracted as described ^107^ using SPLASH®Lipidomix® (Avanti Polar Lipids Inc) as internal standard. Lipid extracts were dissolved in 300 µl chloroform (EMSURE ACS, ISO, Reag. Ph Eur; Supleco):methanol (ULC-MS grade, > 99.97%; Biosolve-Chemicals) (2:1, v/v), 10 µl of the extracts were dried in vacuum and reconstituted in 150 µl chloroform:2-propanol (ULC/MS-CC/SFC grade, >99.95%; Biosolve-Chemicals) (1:1, v/v). Lipids (3 µl) were separated by reverse phase chromatography (Accucore C30 column; 150 mm x 2.1 mm 2.6 µM 150 Å, Thermo Fisher Scientific) using a Vanquish Horizon UHPLC system (Thermo Fisher Scientific) coupled on-line to a Orbitrap Exploris 240 mass spectrometer (Thermo Fisher Scientific) equipped with a HESI source. Lipids were separated at a flow rate of 0.3 ml/min (column temperature 50 °C) and the following gradient: 0-20 min 10 to 80% B, 20-37 min 80 to 95% B,37-41 min 95 to 100% B,41-49 min 100%B, 49.1-57 min 10%B. Eluent A consisted of acetonitrile:water (50:50, v/v, both ULC/MS-CC/SFC grade, Biosolve-Chemicals) and Eluent B of 2-propanol:acetonitrile:water (85:10:5, v/v/v), both containing 5 mM ammonium formate (MS grade, Sigma-Aldrich) and 0.1% formic acid (ULC/MS-CC/SFC grade, Biosolve-Chemicals). Full MS settings were the following: spray voltage – 3500 V, sheath gas – 40 arb units, aux gas – 10 arb units, sweep gas – 1 arb unit, ion transfer tube – 300 °C, vaporizer temperature – 370 °C, EASY-IC run-start, default charge state – 1, resolution at *m/z* 200 – 120.000, scan range – *m/z* 200-1200, normalized AGC target – 100%, maximum injection time – auto, RF lens – 35%. Data-dependent acquisition (DDA) was based on a cycle time (1.3 s) at a resolution of 30.000, isolation window – 1.2 *m/z*, normalized stepped collision energies – 17,27,37%, AGC target – 100%, maximum injection time – 54 ms.

Data analysis was performed using Lipostar2 ^108^ . Briefly, supersample filter considering only lipids with isotopic pattern and with a MS/MS spectrum were kept, detected features were searched against the LIPID MAPS structural database (downloaded 11/2021), and proposed identifications obtained by automatic approval considering 3 and 4 stars manually evaluated. Relative quantities were calculated based on the internal standards and obtained ratios to the extract tissue weight used for extraction.

### Acetyl-CoA measurement

For acetyl-CoA measurement, cultured cells were washed with cold PBS, scraped on ice in cold PBS, centrifuged for 2 min at 1,500 rpm at 4 °C, washed once with 0.1 M ammonium bicarbonate and centrifuged again for 2 min at 1,500 rpm at 4 °C. Cell pellets and tissues were snap-frozen and kept at -80 °C until further analysis. Frozen samples were dissolved in 150 μl 30% methanol in acetonitrile with 100 nM chloropropamide as internal standard. Adipose tissue samples were homogenized in 200 μl 80% isopropanol, 0.5 mm zirconium beads (1/3 volume) were added, samples were homogenized for 15 min at 4 °C and 300 x g in a TissueLyser II (Qiagen) and centrifuged at 13,000 × g for 30 min. Supernatants were collected to be analyzed by LC-MS/MS. For adipose tissue, the supernatant was twice extracted with 1 ml hexane to remove the bulk of di- and triacylglycerols and sterols ^109^. The collected lower fraction was dried and re-dissolved in 200 μl acetonitrile. LC–MS/MS analysis was performed on high performance liquid chromatography (HPLC) system (Agilent 1200) coupled online to G2-S QTof (Waters). For normal phase chromatography Bridge Amide 3.5 μl (2.1×100 mm) columns (Waters) were used. For the normal phase the mobile phase composed of eluent A (95% acetonitrile, 0.1 mM ammonium acetate, and 0.01% NH_4_OH) and eluent B (40% acetonitrile, 0.1 mM ammonium acetate, and 0.01% NH_4_OH) was applied with the following gradient program: Eluent B, from 0% to 100% within 18 min; 100% from 18 to 21 min; 0% from 21 to 26 min. The flow rate was set at 0.3 ml/min. The spray voltage was set at 3.0 kV and the source temperature was set at 120 °C. Nitrogen was used as both cone gas (50 l/h) and desolvation gas (800 l/h), and argon as the collision gas. MSE mode was used in negative ionization polarity. Mass chromatograms and mass spectral data were acquired and processed by MassLynx software (Waters).

### RNAscope

RNAscope was performed in adipose tissue using the RNAscope Multiplex Fluorescent v2 Assay combined with immunofluorescence per manufacturer’s instructions. Snap-frozen SAT and GAT were formalin-fixed paraffin-embedded and cut in 5-µm thick slices. The slides were incubated for 1 h at 60 °C, dehydrated with Roti-clear (CarlRoth) 2 x 5 min and ethanol 2 x 2 min and incubated for 30 min at 60 °C and then 10 min with a H_2_O_2_ solution (ACD) at RT. Then, they were washed with distilled water, incubated for 15 min with CO-Detection Target Retrieval (ACD) using a pressure cook, rinsed with distilled water and washed 3 x 5 min with PBS-T. Antibodies anti-PDGFRα (1:200, AF1062, R&D Systems), anti-perilipin (1:100, GP29, Progen) and anti-caveolin (1:150) diluted in CO-Detection Buffer (ACD) were added overnight at 4 °C. Then, slides were washed 2 x 2 min with PBS-T and fixed with neutral buffered formalin (Morphisto GmbH) 30 min at RT, washed 2 x 2 min with PBS-T and incubated with Protease Plus (ACD) 15 min at 40 °C using the HybEZ II hybridization system oven (ACD). After washing with distilled water, the samples were let to hybridize for 2 h at 40 °C in the HybEZ II oven with the Probe-Mm-*Azin2* (569871, Bio-techne). The slides were then hybridized with Amplifier (AMP)1, AMP2 and AMP3 for 30 min at 40 °C in the HybEZ II oven with 2 x 2 min washes with ACD Wash Buffer in between hybridizations. Afterwards, slides were sequentially incubated with RNAscope Multiplex FL v2 HRP-C1, OPAL 570 (Akoya Biosciences) and RNAscope Multiplex FL v2 HRP Blocker, all for 15 min at 40 °C in the HybEZ II oven with washes with ACD Wash Buffer in between incubations. Then, samples were incubated for 30 min at RT with donkey anti-goat IgG, Alexa Fluor™ 647 (A21447, Thermo Fisher Scientific) and goat anti-guinea pig IgG H&L Alexa Fluor® 488 (ab150185, Abcam) diluted in CO-Detection AB Diluent (ACD). Finally, they were washed 2 x 2 min with PBS-T, DAPI (1:10,000, Sigma-Aldrich) and then TrueBlack Lipofuscin Quencher (Thermo Fisher Scientific) were added for 30 sec, and slides were mounted with Prolong Gold Antifade Mountant (Thermo Fisher Scientific). Adipose tissue sections were imaged with a ZeissLSM880 (Zeiss).

### Immunofluorescence in adipose tissue

Immunofluerescent stainings were performed as previously described ^91, 52^. Snap-frozen SAT and GAT were formalin-fixed, paraffin-embedded and cut in 5-µm thick slices. They were dehydrated 3 x 5 min with Roticlear (Carlroth), sequential 5-min washings with ethanol 100, 95, 80 and 70% ethanol solutions, and a 5-min wash with PBS. Antigen retrieval was performed for 10 min with Target Retrieval Solution, Citrate pH 6 (Agilent) using a pressure cooker, followed by permeabilization for 10 min in 0.1% Triton X-100 in PBS, 3 x 5 min washes with PBS and blocking 2 h with Protein Block, Serum-Free (Agilent). Then, samples were incubated overnight at 4 °C with following primary antibodies: anti-Arginase 1 (1:50, BD Biosciences, 610709), anti-SAT1 (1:100, Novus Biologicals, B110-41622), anti-ODC1 (1:100, Biorbyt, orb228847), anti-IL4Rα (1:50, Biorbyt, orb337363), anti-PDGFRα (1:200, R&D Systems, AF1062), anti-Perilipin (1:100, Progen, GP29) or anti-Caveolin (1:150 ab192869 Abcam), all diluted in Antibody Diluent (Agilent). After samples were washed 3 x 5 min with PBS they were incubated for 1.5 h with following secondary antibodies all diluted 1:350 in Antibody Diluent (Agilent) at RT: donkey anti-mouse IgG, Alexa Fluor™ 647 (A-31571, Thermo Fisher Scientific), donkey anti-rabbit, Alexa Fluor™ 647 (A-31573, Thermo Fisher Scientific), donkey anti-goat IgG, Alexa Fluor™ 555 (A-21432, Thermo Fisher Scientific), goat anti-guinea pig IgG H&L Alexa Fluor® 488 (ab150185, Abcam) or donkey anti-rabbit Alexa Fluor 488 (A21206 Invitrogen). Samples were then washed 3 x 5 min with PBS, incubated for 30 sec with TrueBlack® Lipofuscin Autofluorescence Quencher (Biotium), washed 3 x 5 min with PBS, stained with DAPI (1:10,000, Sigma-Aldrich) and mounted with Fluoromount-G™ Mounting Medium (Thermo Fisher Scientific). Adipose tissue sections were imaged with a Zeiss LSM880.

### Histology

Fresh tissues were excised, snap-frozen, fixed in 4% paraformaldehyde and embedded in paraffin. For adipocyte size analysis, sections were stained with hematoxylin and eosin (H&E). F4/80 staining, antigen retrieval was performed with citrate buffer pH 6.0, and heating for 10 min, followed by 10 min peroxidase blocking. Following steps were performed with the VECTASTAIN® Elite® ABC Universal PLUS Kit (Vector Laboratories). Samples were incubated overnight at 4 °C with anti-F4/80 (NB:600-404 Novus Biologicals) diluted at 1/125 in antibody diluent (Dako). After washing 3 x 5 min with PBS-T they were incubated for 1 h with anti-rat secondary IgG antibody at RT (PI-9400-1 Vector Laboratories). To determine fibrosis tissues sections were stained for one hour with Picrosirius red staining solution followed by 5x washes with 1% acetic acid. Images were acquired at 10x magnification using a ZEISS OBSERVER Z.1 - widefield microscope with apotome (Zeiss). Images of Picrosirius red staining were acquired with polarized light. Plugin macros in FIJI software were used for quantification of adipocyte size and Picrosirius red staining^110^.

### Nile red staining

Liver tissues were embedded in OCT. Sections were fixed for 30 min in 4 % PFA, washed 3 x 5 min with PBS and stained for 1 h at 37°C with the Nile Red Staining Kit (ab228553 Abcam), followed by DAPI staining and mounting. Images were acquired with an Axio Observer Z1/7 inverted microscope with Apotome mode (Zeiss) at 10 x magnification and the ZEN 3.2 blue edition software.

### Oil Red O staining

Differentiated APs were washed twice with PBS, fixed with 10% formalin for one hour, rinsed with distilled water, and stained for 1 h with Oil Red O dye in 60% isopropanol. Then, they were thoroughly rinsed with distilled water and images were acquired at bright field with an Axio Observer Z1/7 inverted microscope with Apotome mode (Zeiss) and the ZEN 3.2 blue edition software. At least 5 view-fields were imaged per sample. The stain was extracted through incubation with 100% isopropanol for 5 min. Absorbance was measured at 492 nm using a microplate reader (Biotek).

### Immunofluorescence in cultured cells

Differentiated APs were fixed for 20 min with 4% paraformaldehyde, washed 3×5 min with PBS, permeabilized for 10 min with 0.1 % Triton X-100, washed again 3×5 min with PBS and blocked with blocking buffer (DAKO). Samples were then incubated overnight at 4 °C with anti-PPARG (1:100; #2443, Cell Signaling). After 3×5 min with PBS washes they were incubated for 1.5 h at RT with anti-rabbit Alexa Fluor 555 IgG (A-31572 ThermoFisher, Scientific) diluted 1:350 in antibody diluent (DAKO). Finally, they were washed 3×5 min with PBS, stained with DAPI and mounted. Images were acquired with an Axio Observer Z1/7 inverted microscope with Apotome mode (Zeiss) and the ZEN 3.2 blue edition software.

For CDKN2A and CDKN1A staining, cells were fixed for 10 min with 4% paraformaldehyde at RT and 3x washed with PBS. Then, they were permeabilized for 6 min with 0.1% Triton X-100 in PBS, washed 3×5 min washes with PBS and blocked for 1 h with Protein Block, Serum-Free (Agilent). Then, they were incubated overnight at 4 °C with anti-CDKN2A (1:100, ab211542, Abcam) or anti-CDKN1A (1:200, ab188224, Abcam) diluted in Antibody Diluent (Agilent). Afterwards, they were washed 3×5 min with PBS and incubated for 2 h with donkey anti-rabbit-Alexa Fluor™ 555 (1:350, A-31572, Thermo Fisher Scientific). Cells were washed 3×5 min with PBS, stained with DAPI (1:10,000, Sigma-Aldrich) and mounted with Fluoromount-G™ Mounting Medium (Thermo Fisher Scientific). Slides were imaged with a Zeiss LSM880.

### Senescence β-Galactosidase Staining

Differentiated APs were stained with the Senescence β-Galactosidase Staining Kit (#9860 Cell Signaling) following manufacturer’s instructions. Images were taken with an Axio Observer Z1/7 inverted microscope with Apotome mode (Zeiss) and the ZEN 3.2 blue edition software. At least 5 view-fields were imaged per sample.

### Image acquisition and analysis

RNAscope and immunofluorescence images were acquired on a Zeiss LSM 880 inverted confocal microscope (Zeiss), illuminated with laser lines at 405 nm, 488 nm, 561 nm and 633 nm, and detected by two photomultiplier tube detectors. EC Plan-Neofluoar objectives with 20x and 40x magnification, 0.8 or 1.30 numerical aperture and M27 thread, working with an oil immersion medium Immersol 518 F, were used. Laser power, photomultiplier gain and pinhole size were set for each antibody individually and kept constant for all image acquisitions. Images were acquired with the ZEN 2.3 black edition software. Image processing was performed with Imaris 9/10 (Bitplane AG) with Labkit Plugin. Channels were Median filtered (3×3×1) and DAPI/Perilipin channels were rendered using the Imaris “Surface” pipelines, to outline the nucleus and adipocyte cell body, respectively. Channels were contrast adapted by linear stretch using the same limits across a sample set to make intensities comparable. Maximum intensity projections (MIP) were computed and “Snapshot” was taken. Video files were created by the “Animation” function of Imaris. Images are shown in pseudocolor; the display color of the channels was set as to optimize clarity of merged images.

Microscopic images of H&E stainings were acquired with an Axio Observer Z1/7 inverted microscope (Zeiss), on a Plan-Apochromat objective with 10x magnification, 0.30 numerical aperture and M27 thread. For each mouse at least 8 view-fields were imaged per tissue section. Images were acquired with the ZEN 3.2 blue edition software and processed and quantified with the Fiji software.

### FACS analysis

APs and ATMs were analysed or sorted by FACS from the SVF of the adipose tissue using following antibodies: anti-CD140a (PDGFRα)-APC (1:100, 562777, BD Biosciences), anti-LY6A-PECy7 (1:100, 558162, BD Biosciences), anti-CD31-PE (1:100, 553373, BD Biosciences) and anti-CD45-AlexaFluor488 (1:100, 103122, BioLegend), anti-CD45-PE (1:100, 553081, BD Biosciences), anti-CD11b-PerCP (1:100, 101230, BioLegend), anti-F4/80-PECy7 (1:100, 25480182, eBioscience), anti-CD11c-APC (1:100, 117310, BioLegend) and anti-CD9-FITC (1:100, 11009182, Invitrogen). For Ki67 staining, cells were permeabilized with the Fix/Perm Buffer Set (Biolegend) and incubated with anti-Ki67-FITC (1:2,000, 11-5698-82, Thermo Fisher Scientific) for 45 min at 4 °C. FACS was carried with a BD FACSCanto II (BD Biosciences) and analyzed with the BD FACSDiva Version 6.1.3 software (BD Biosciences).

### Triglyceride measurement

Triglyceride content quantification in the liver was performed with a kit (Abcam), as previously described ^30^. Briefly, 100 mg liver tissue were homogenized in 1 ml of 5% Triton X-100, heated at 95°C and cooled to room temperature twice. Thereafter, samples were centrifuged, and triglyceride content in the supernatant was quantified following manufacturer’s instructions.

### Western blot

Cultured 3T3-L1 cells were lysed in lysis buffer containing 1% SDS, 50 mM Tris pH 7.4, 1 mM sodium orthovanadate, protease cOmplete™ Protease Inhibitor Cocktail inhibitors (Merck) and DNase (benzonase, Sigma). Cell lysates were centrifuged at 16,000 g for 5 min at 4 °C, supernatants were collected and total protein concentration was measured using Pierce BCA Protein Assay Kit (Thermo Fisher Scientific). 40 μg of protein were loaded onto polyacrylamide gels and proteins were transferred on a nitrocellulose membrane by wet transfer. Membranes were blocked in 5% non-fat milk or 5% BSA TBS-T (0.15 M NaCl, 2.7 mM KCl, 24.8 mM Tris-base, 0.1% Tween-20). Membranes were incubated overnight at 4 °C with an anti-DDK (1:1,000, TA50011-100, Origene). Membranes were then incubated for 2 h at RT with goat anti-mouse IgG HRP-conjugated (1:3,000; Jackson ImmunoResearch) diluted in 5% non-fat milk in TBS-T. Membranes were revealed using SuperSignal West Pico Chemiluminescent Substrate (Thermo Fisher Scientific) and a luminescent image analyzer, LAS-3000 (Fujifilm).

### qPCR

RNA was isolated from frozen tissues with the Trizol Reagent (Thermo Fisher Scientific) after mechanical tissue disruption or from cells with the Nucleospin RNA kit (Macherey-Nagel) according to manufacturer’s instructions. RNA isolated with Trizol was subsequently treated with DNase (Thermo Fisher Scientific). Reverse transcription was performed with the iScript cDNA Synthesis kit (Biorad) and cDNA was analysed by qPCR using the SsoFast Eva Green Supermix (Bio-Rad), a CFX384 real-time System C1000 Thermal Cycler (Bio-Rad) and the Bio-Rad CFX Manager 3.1 software. *18S* expression was used as an internal control for normalization. Primers used are shown in Table S1.

### Statistical methods

Statistical analysis and data plotting were done with the GraphPrism 7.04 software. Statistical tests used are described in each figure legend, p < 0.05 or adjp < 0.05 was set as a significance level.

### Graphical design

Schemes were designed with Biorender.com.

## Supplementary figure legends

**Figure S1.**
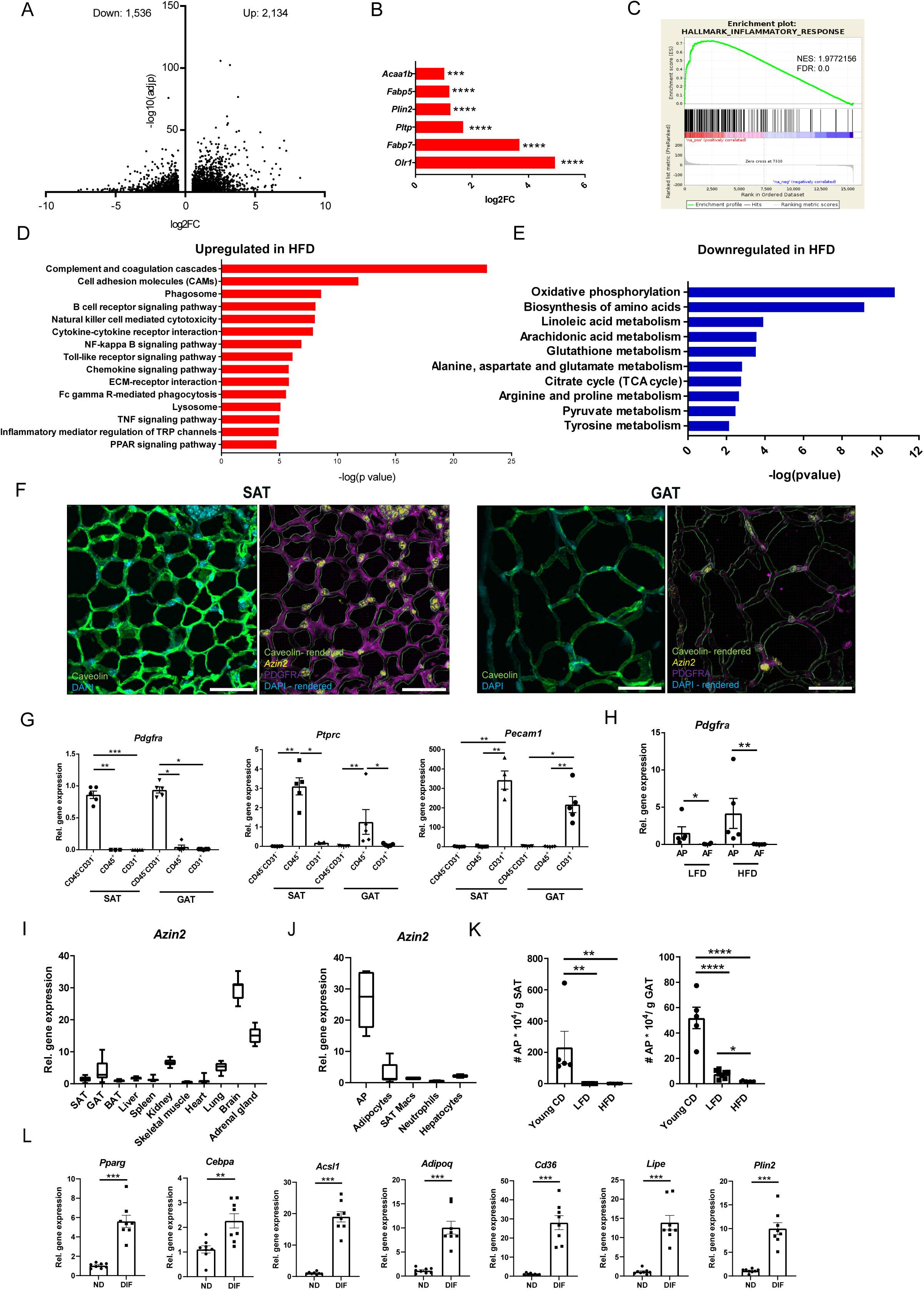
Polyamine metabolism is downregulated in the adipose tissue with obesity, related to Figure 1. **A-D.** RNA-seq in SAT of HFD compared to LFD mice (n=4 mice per group). Volcano plot of differentially expressed genes with log2FC > 0.5 and <-0.5 **(A)**. Differentially expressed adipogenic genes in SAT of HFD compared to LFD mice **(B)**. GSEA analysis for inflammatory response-related gene set **(C)**. Upregulated KEGG signaling pathways **(D)** and downregulated KEGG metabolic pathways **(E)** in SAT of HFD mice, as shown by EGSEA analysis (adjp<0.05). **F.** RNAScope for *Azin2* and immunofluorescence for PDGFRα and Caveolin in SAT and GAT of 10 week-old mice fed a chow diet (CD). Scale bar, 50 μm. Representative images from 2 mice, 3 levels per tissue are shown. **G.** *Pdgfra*, *Ptprc* (*Cd45*) and *Pecam1* (*Cd31*) expression in CD45^+^, CD31^+^ and CD45^-^CD31^-^ SVF populations isolated from SAT and GAT of mice fed for 20 weeks a LFD (n=5 mice per group). **H.** *Pdgfra* expression in SAT APs and AF of mice fed for 20 weeks a LFD or a HFD (n=4-5 mice per group). **I-J.** *Azin2* expression in different mouse tissues (n=8 mice) (**I**) and cell types (n=2-6 mice) (**J**). **K.** Number of APs (PDGFRA^+^Sca1^+^CD31^-^CD45^-^) per g of adipose tissue assessed by FACS in SAT and GAT of young (CD-fed), LFD- and HFD-fed wt mice (n=5-8 mice per group). **L.** *Pparg*, *Cebpa*, *Acsl1*, *Adipoq*, *Cd36*, *Lipe* and *Plin2* expression in non-differentiated and differentiated SAT APs (n=8 mice per group). Data in **G-L** are shown as mean±SEM, Kruskal-Wallis (**G,K**) and Mann-Whitney U (**H, L**) tests were used for statistical analysis. *p < 0.05, **p < 0.01, ***p < 0.001. NES: Normalized enrichment score; FDR: False discovery rate; SAT: subcutaneous adipose tissue; GAT: gonadal adipose tissue; BAT: brown adipose tissue; CD: chow diet; LFD: Low fat diet; HFD: High fat diet; AP: adipocyte progenitors; AF: adipocyte fraction; Macs: Macrophages

**Figure S2.**
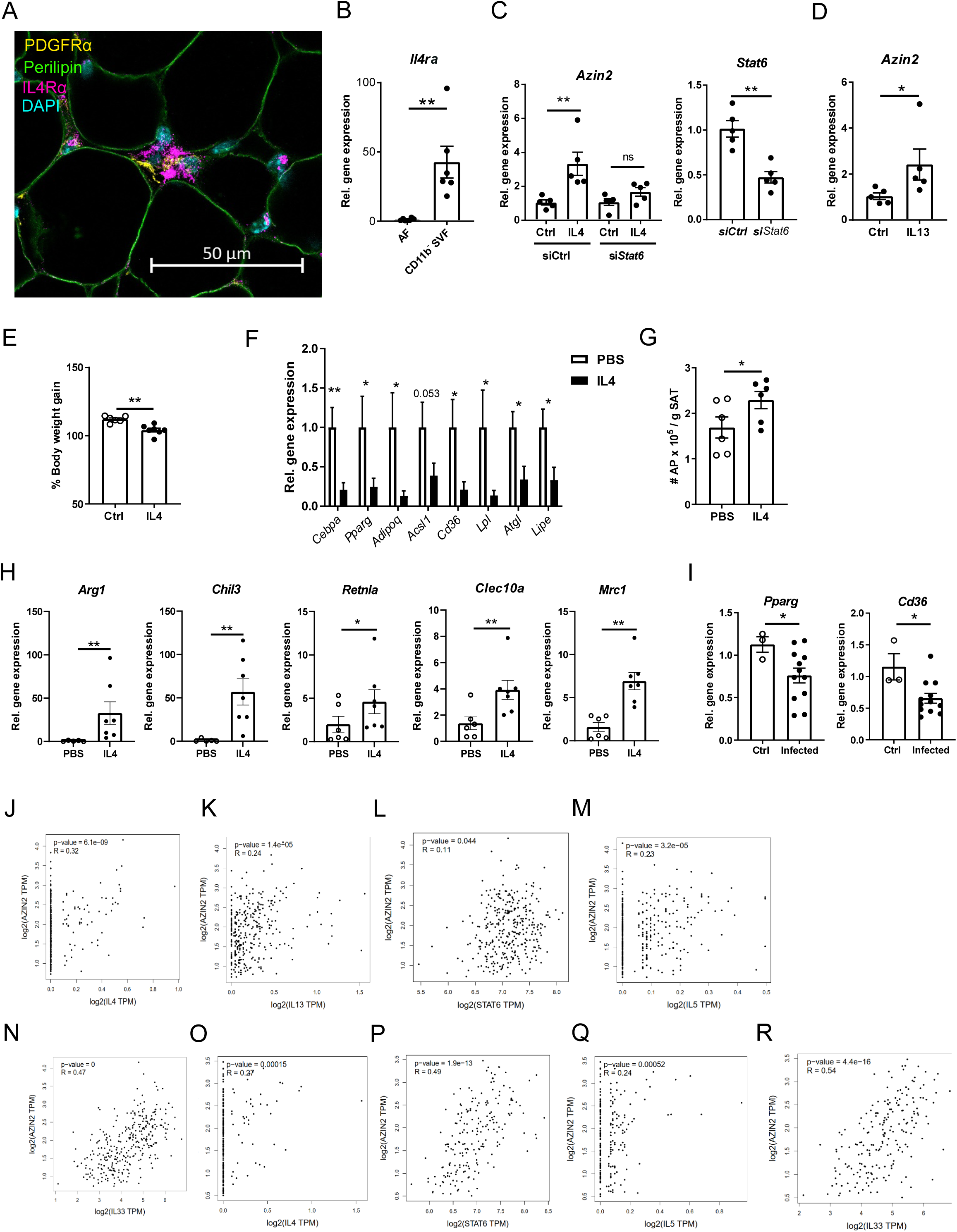
IL4 upregulates *Azin2* expression in APs, related to Figure 2. **A.** Immunofluoeresence image of SAT stained for IL4Rα (magenta), PDGFRα (yellow), Perilipin (green) and DAPI (cyan). Scale bar, 50 μm **B.** *IL4ra* expression in the AF and CD11b^-^ SVF cells of SAT of HFD mice (n=6). **C.** *Azin2* and *Stat6* expression in non-differentiated SAT APs transfected with si*Stat6* or siCtrl for 24 h and treated or not for 4 h with IL4 (20 ng/ml) (n=5). **D.** *Azin2* expression in non-differentiated SAT APs treated for 4 h with IL13 (20 ng/ml) (n=5). **E-H**. Wt mice were fed for 6 weeks a HFD and treated every 2 days with PBS or IL4 + a-IL4 during the last 2 feeding weeks. The body weight gain (**E**), adipogenic gene expression (**F**) and AP number in SAT (**G**) are shown (n=6-7 mice per group). **H.** *Arg1*, *Chil3*, *Retnla*, *Clec10a* and *Mrc1* expression in SAT SVF of PBS and IL4-treated mice (n=6-7 mice per group). **I.** *Pparg* and *Cd36* expression in GAT of mice infected with *N. brasiliensis* or control mice 9 days post-infection (n=4-9 mice per group). **J-R.** Correlation of *AZIN2* expression with *IL4, IL13, STAT6, IL5,* and *IL33* in human subcutaneous (**J-M**) and visceral (**N-R**) adipose tissue based on the GEPIA2 database (http://gepia2.cancer-pku.cn/#index). Data in **C-I** are shown as mean±SEM. Gene expression in **B-D**, **F, H** and **I** was determined by qPCR using *18S* as a housekeeping gene. Mann-Whitney U test was used for statistical analysis. *p < 0.05, **p < 0.01

**Figure S3.**
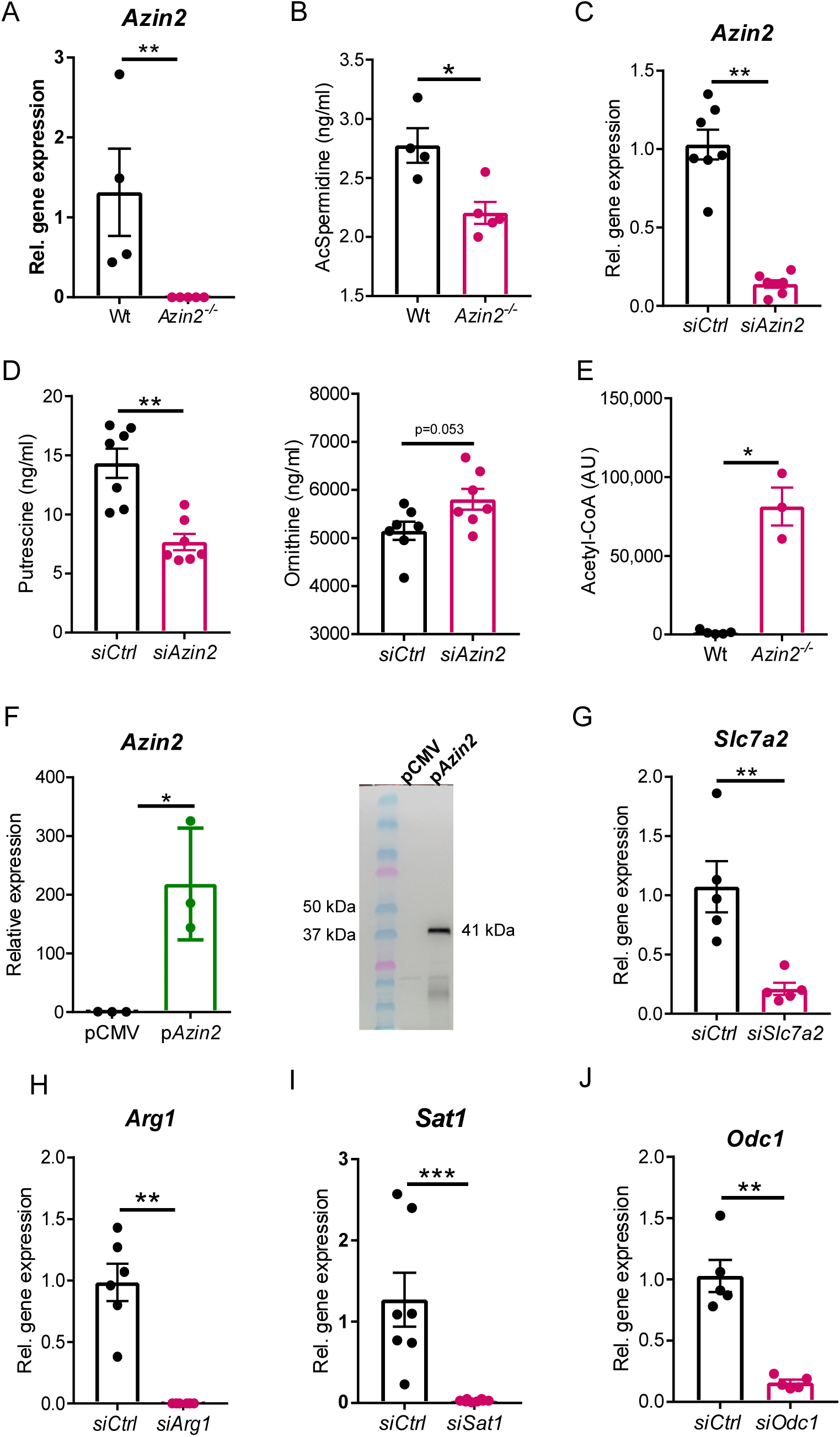
AZIN2 promotes polyamine acetylation and negatively regulates acetyl-CoA levels in APs, related to Figure 3. **A.** *Azin2* expression in differentiated wt and *Azin2^-/-^* SAT APs (n=4-5 mice per group). **B.** N1-acetylspermidine concentration in the cell culture supernatant of in vitro differentiated wt and *Azin2^-/-^* SAT APs (n=4-5 mice per group). **C.** *Azin2* expression in SAT APs transfected 3 times over one week of differentiation (every 2-3 days), with si*Azin2* or control siRNA (n=5-7 mice per group). **D.** Putrescine and ornithine levels in cell culture supernatants of SAT APs 24 h after transfection with si*Azin2* or control siRNA or (n=7 mice per group). **E.** Acetyl-CoA levels in differentiated wt and *Azin2^-/-^* SAT APs (n=3-5 mice per group). **F.** *Azin2* mRNA expression (left) and western blot for AZIN2-DDK (right) in 3T3-L1 preadipocytes transfected for 24 h with a plasmid overexpressing AZIN2-DDK or control pCMV plasmid (n=3 for left panel). **G-I.** *Slc7a2*, *Arg1*, and *Sat1* expression in SAT APs transfected 3 times over one week of differentiation with the respective siRNA (n=5 for **G**, n=6 for **H**, n=7-8 for **I**). **J.** *Odc1* expression in SAT APs 24 h after transfection with si*Odc1* or control siRNA (n=5 mice per group). Gene expression was determined by qPCR using *18S* as housekeeping gene. Polyamine levels were determined by LC-MS/MS. Data are shown as mean±SEM. Mann-Whitney U test was used for statistical analysis. *p < 0.05, **p < 0.01, ***p < 0.001

**Figure S4.**
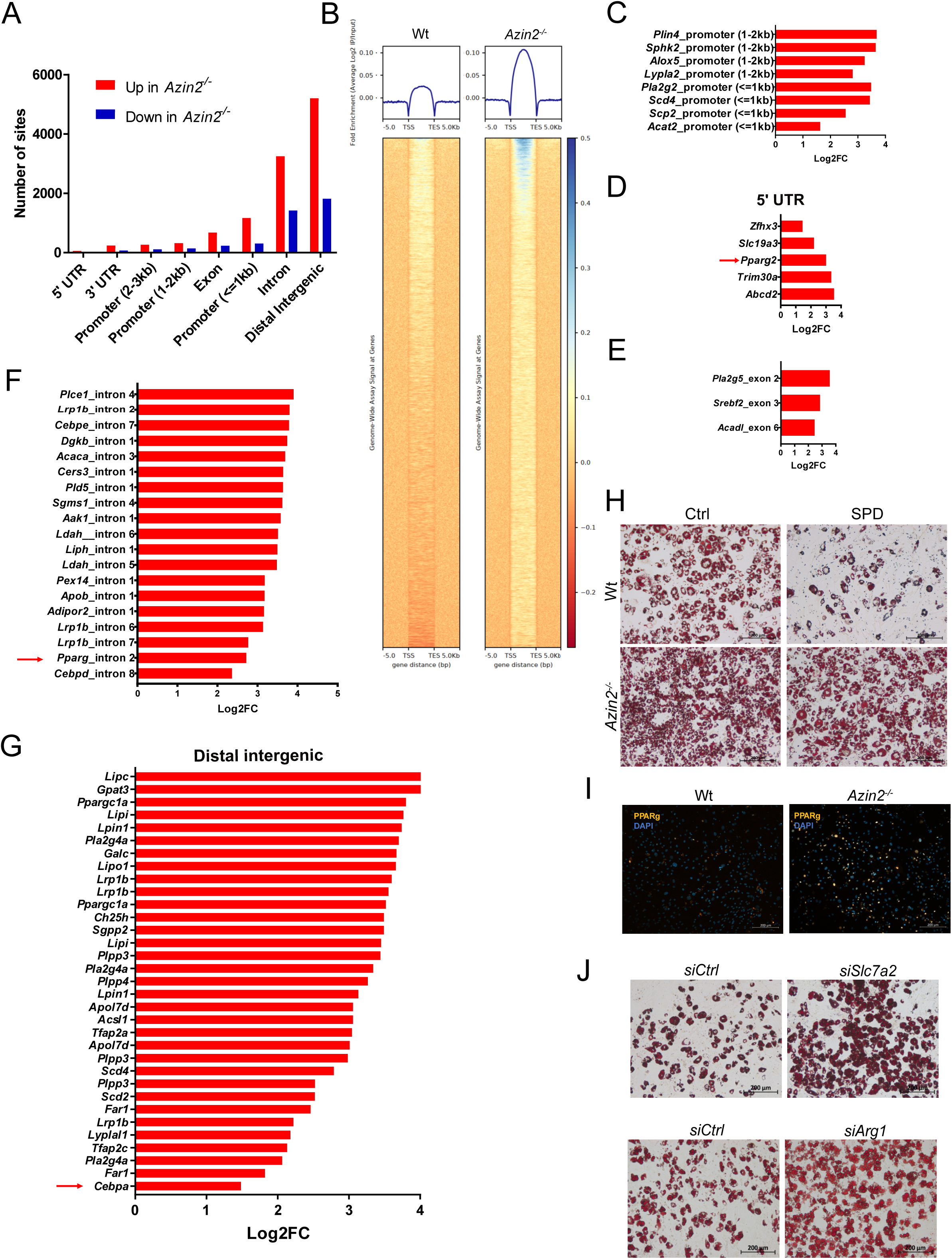
AZIN2 inhibits adipogenesis through regulation of histone acetylation, related to Figure 4. **A.** Number of sites with enriched H3K27ac marks in different DNA regions up- or downregulated in *Azin2^-/-^* compared to wt differentiated SAT APs. **B.** Genome-wide average log2FC enrichment of H3K27ac marks in regions between transcriptional start sites (TSS) and transcriptional end sites (TES) for representative wt and *Azin2^-/-^* SAT AP samples. **C-G.** Genes with H3K27ac enrichment in the promoter (1-2 kb and <1kb) (**C**), 5’ UTR region (**D**), exons (**E**), introns (**F**) and distal intergenic regions (**G**) in *Azin2^-/-^* versus wt SAT APs. The *Pparg* gene is depicted with an arrow. **A-G**: n=3 pools of APs from 2 mice per group. **H.** Images of Oil-Red O staining in differentiated wt and *Azin2^-/-^* SAT APs treated during differentiation with spermidine (10 μM). Representative images of cells from 1 out of 8 mice is shown per condition, scale bar: 200 μm. **I.** Images of immunofluorescence for PPARg in differentiated wt and *Azin2^-/-^* SAT APs. Representative images of cells from 1 out of 8 mice is shown per condition, scale bar: 200 μm. **J.** Images of Oil-Red O staining in SAT APs transfected 3 times over one week of differentiation with siRNA against *Slc7a2*, *Arg1* or control siRNA. Representative images of cells from 1 out of 5-8 mice per condition are shown, scale bar: 200 μm.

**Figure S5.**
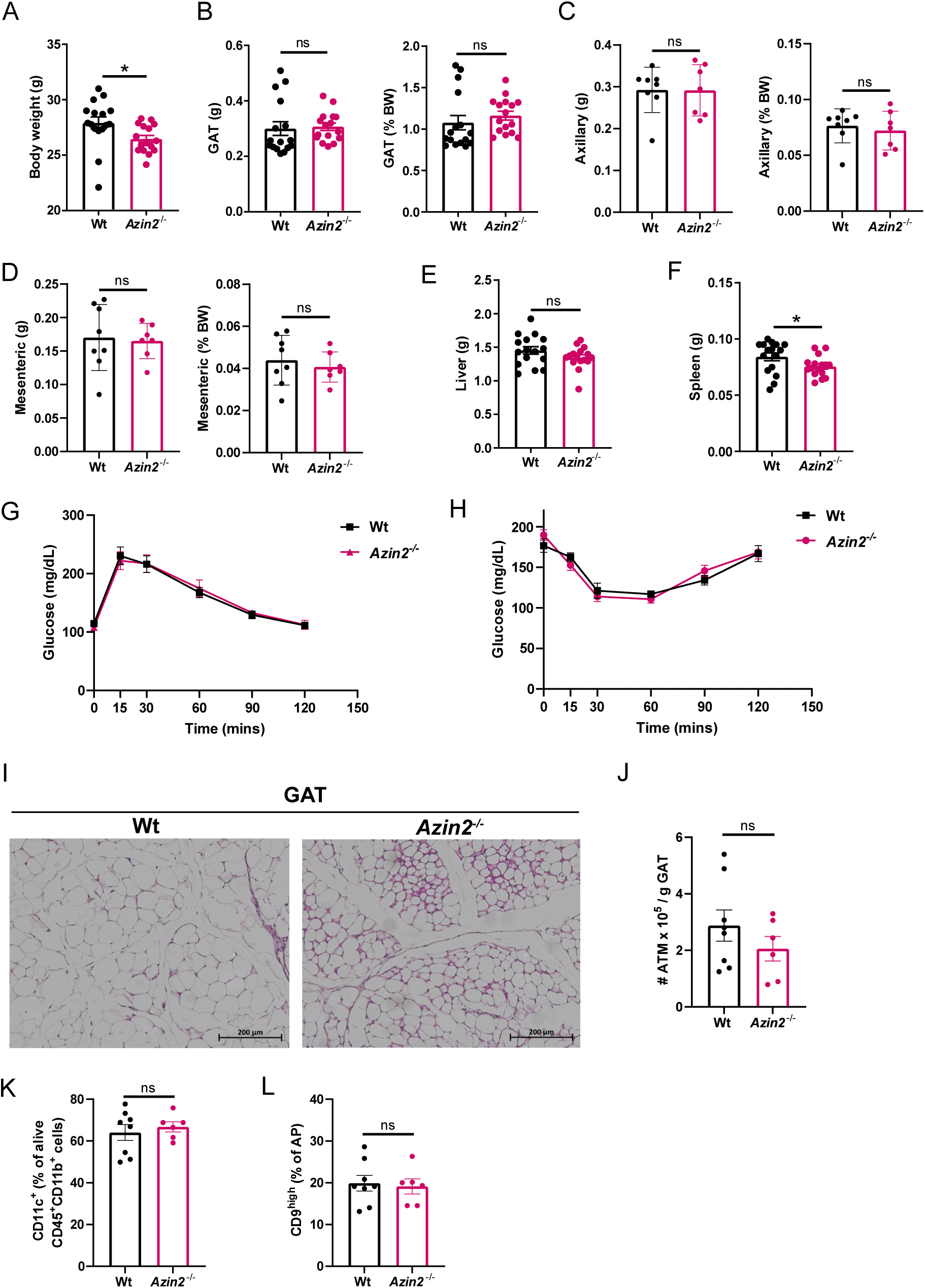
In the lean state, GAT in AZIN2 deficient mice is composed of more and smaller adipocytes compared to wt mice, related to Figure 5. **A-F.** Body weight (**A**), GAT weight (**B**), axillary adipose tissue weight (**C**), mesenteric adipose tissue weight (**D**), liver weight (**E**) and spleen weight (**F**) in 10 weeks-old chow diet-fed wt and *Azin2^-/-^* mice (n=16 mice per group for **A-C, E,F** and n=6-8 mice per group for **C** and **D**). **G,H.** Glucose tolerance test (GTT) (**G**) and insulin tolerance test (ITT) (**H**) in 10 weeks-old chow diet-fed wt and *Azin2^-/-^* mice (n=6-8 mice per group). **I.** Representative H&E images of GAT in 10 weeks-old chow diet-fed wt and *Azin2^-/-^* mice, scale bar: 200 μm (n= 3 mice per group). **J.** Number of ATMs (CD45^+^CD11b^+^F4/80^+^) per g of GAT assessed by FACS in wt and *Azin2^-/-^* mice (n=6-8 mice per group). **K.** Percentage of CD11c^+^ ATMs (CD45^+^CD11b^+^) in GAT of wt and *Azin2^-/-^* mice assessed by FACS (n=6-8 mice per group). **L.** Percentage of CD9^high^ APs (PDGFRα^+^LY6A^+^CD31^-^CD45^-^) in GAT of wt and *Azin2^-/-^* mice assessed by FACS (n=6-8 mice per group). Data in **A-H** and **J-L** are shown as mean±SEM. Mann-Whitney U test was used for statistical analysis. *p < 0.05, ns: not significant.

**Figure S6.**
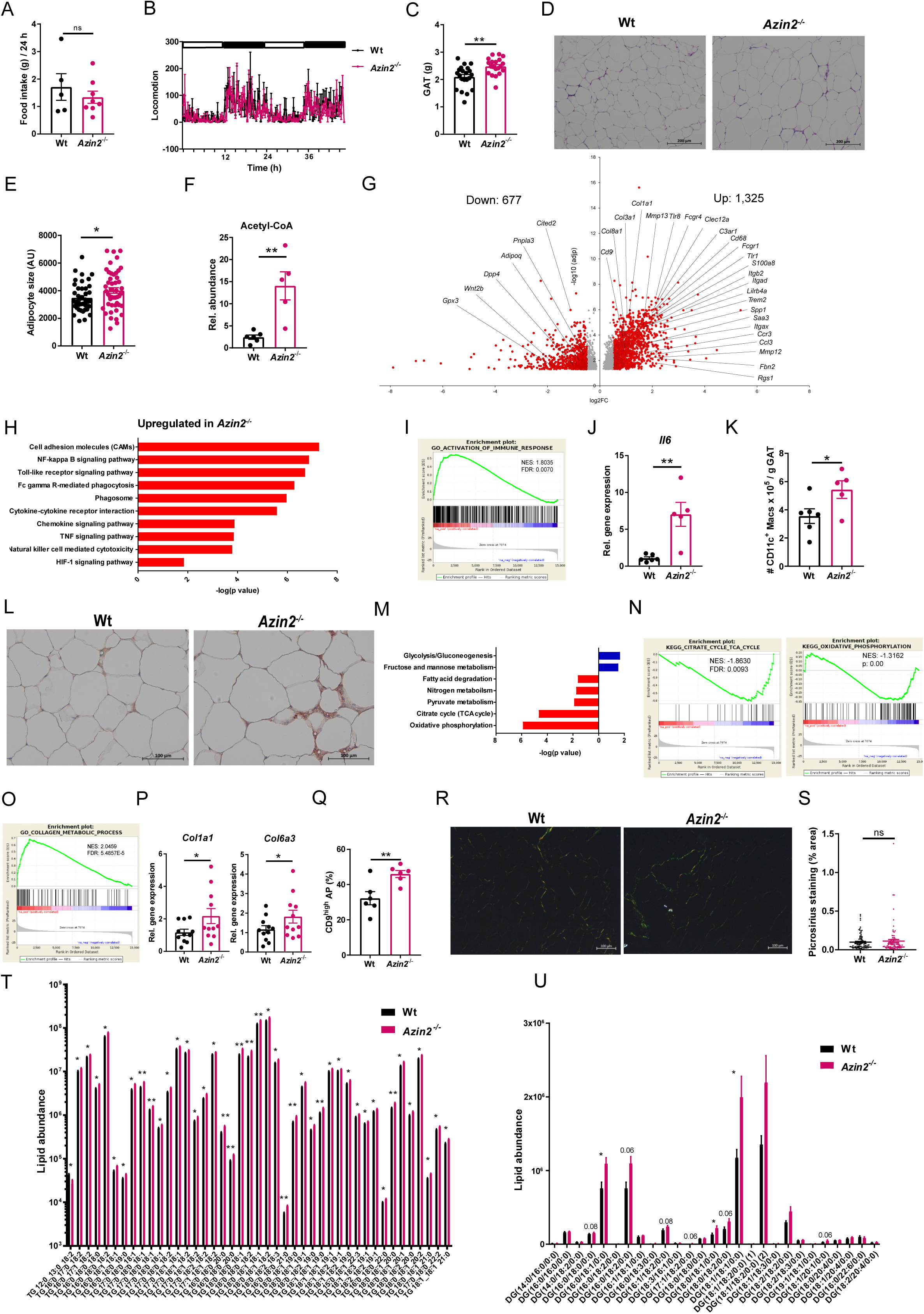
AZIN2 deficiency leads to increased obesity and adipose tissue dysfunction, related to Figure 6. **A,B.** Food uptake and locomotion measured in metabolic cages in wt and *Azin2^-/-^* mice fed for 6 weeks a HFD (n=5-8 for **A** and 7-9 for **B** mice per group). **C.** GAT weight of wt and *Azin2^-/-^* mice fed for 8 weeks a HFD (n=20-21 mice per group). **D.** Representative images of H&E staining of GAT from wt and *Azin2^-/-^* mice fed for 8 weeks a HFD (one image is shown from one out of 10-11 mice), scale bar: 200 μm. **E.** Adipocyte size quantification in GAT of wt and *Azin2^-/-^* mice fed for 8 weeks a HFD (n=6 mice per group, 9-10 sections per mouse were imaged, each dot represents the size of the sliced area of one adipocyte). **F.** Relative abundance of acetyl-CoA levels in GAT of wt and *Azin2^-/-^* mice fed for 8 weeks a HFD presented as peak intensity assessed by LC-MS/MS (n=4-6 mice per group). **G-I, M-O.** Bulk RNA-seq in GAT of wt and *Azin2^-/-^* mice fed for 8 weeks a HFD (n=4 mice per group): Volcano plot of differentially expressed genes with log2FC > 0.5, < -0.5 (**G**), EGSEA analysis for upregulated KEGG signaling pathways (padj<0.05) in *Azin2^-/-^* mice (**H**), GSEA analysis for GO_Activation of Immune Response (**I**). EGSEA pathway analysis showing up-(blue) and downregulated (red) KEGG metabolic pathways (padj<0.05) in *Azin2^-/-^* mice (**M**), GSEA analysis for KEGG_Citrate cycle and KEGG Oxidative Phosphorylation (**N**) and GO_Collagen Metabolic Process (**O**). **J.** *Il6* expression in GAT SVF of wt and *Azin2^-/-^* mice fed for 8 weeks a HFD (n=5-6 mice per group). **K.** Number of CD11c^+^ macrophages (CD45^+^CD11b^+^) in GAT of wt and *Azin2^-/-^* mice fed for 8 weeks a HFD assessed by FACS (n=5-6 mice per group). **L.** F4/80 staining in GAT of wt and *Azin2^-/-^* mice fed for 8 weeks a HFD, scale bar: 200 μm (representative images from one out of 3 mice per group). **P** *Col1a1* and *Col6a3* expression in GAT of wt and *Azin2^-/-^* mice fed for 8 weeks a HFD (n=11 mice per group). **Q.** Percentage of CD9^high^ GAT APs (PDGFRα^+^LY6A^+^CD31^-^CD45^-^) in wt and *Azin2^-/-^* mice fed for 8 weeks a HFD (n=6 mice per group). **R,S.** Picrosirius staining (**R**) and quantification (as % of area) (**S**) in GAT of wt and *Azin2^-/-^* mice fed for 8 weeks a HFD (**R**: representative images from one out of 10-11 mice per group**, S:** n=10-11 mice per group, 8-10 images per mouse) **T,U.** Relative abundance of TG species (**T**) and DG species (**U**) in SAT of wt and *Azin2^-/-^* mice fed for 8 weeks a HFD shown as peak intensity determined by LC-MS/MS, in **T** only differentially regulated TG species are shown (n=5-8 mice per group). Gene expression in **J** and **P** was assessed by qPCR using *18S* as a housekeeping gene. Data in **A-C, E, F, J, K, P, Q, S, T** and **U** are shown as mean±SEM. Mann-Whitney U test (for **A-C,F,J,K,P,Q**), t-test (**E,S**) and multiple t-test (**T,U**) were used for statistical analysis. *p < 0.05, **p < 0.01, ns: not significant. NES: Normalized enrichment score; FDR: False discovery rate

**Figure S7.**
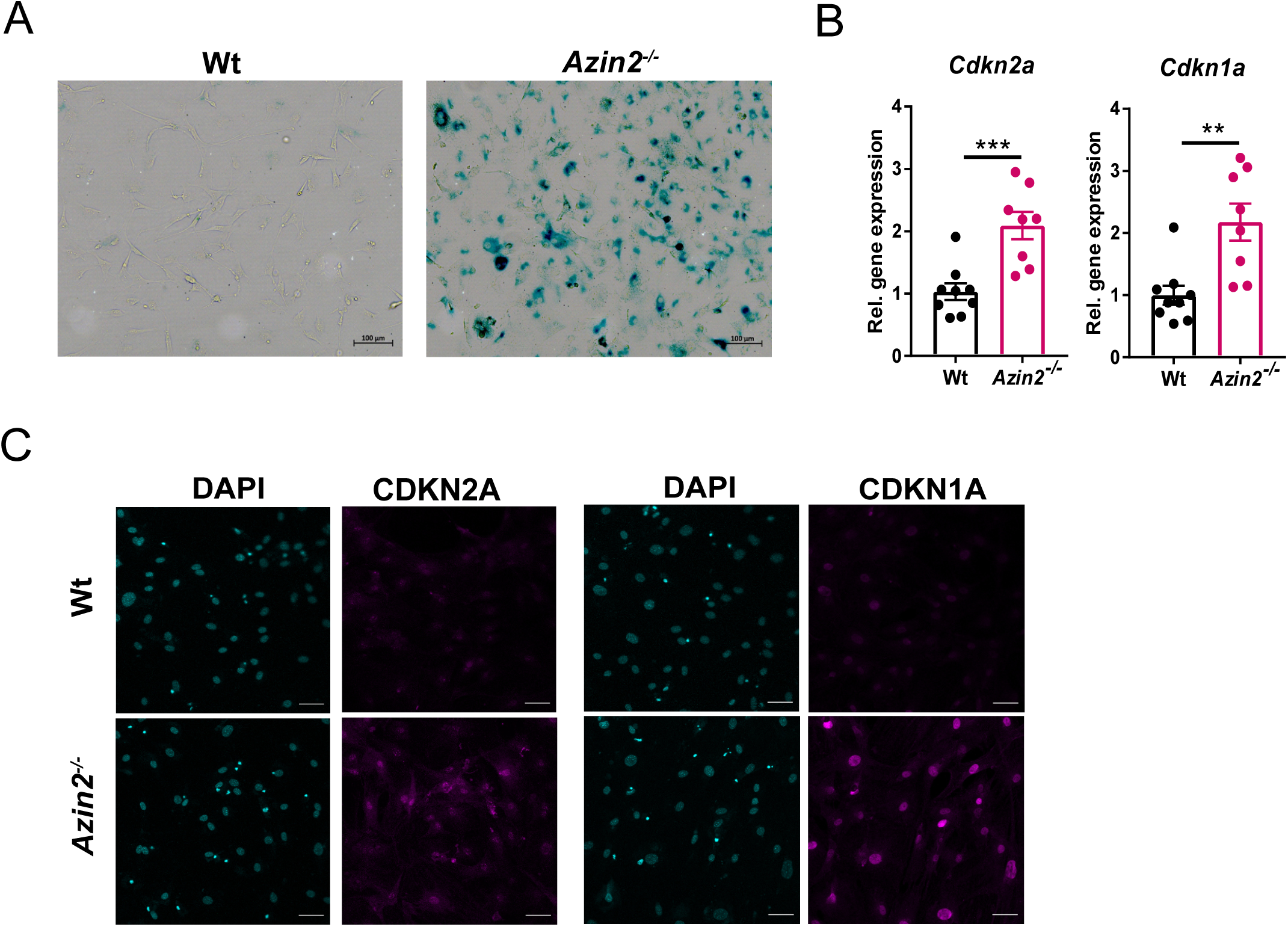
AZIN2 deficiency leads to increased AP senescence, related to Figure 6. Wt and *Azin2^-/-^* SAT APs were differentiated for one week and cultured for another week to induce senescence. **A.** Senescence was assessed by β-Galactosidase staining (one representative image from 1 out of 5 mice per group is shown). **B.** *Cdkn2a* (*p16*) and *Cdkn1a* (*p21*) mRNA expression was assessed by qPCR using *18S* as a housekeeping gene (n=8-9 mice per group). **C.** Immunofluorescence staining against CDKN2A or CDKN1A (magenta) and DAPI staining (cyan); Scale bar, 50 μm; (one representative image from 1 out of 3-4 mice per group is shown). Data in **A** are shown as mean±SEM. Mann-Whitney U test was performed **p < 0.01, ***p < 0.001.

**Figure S8.**
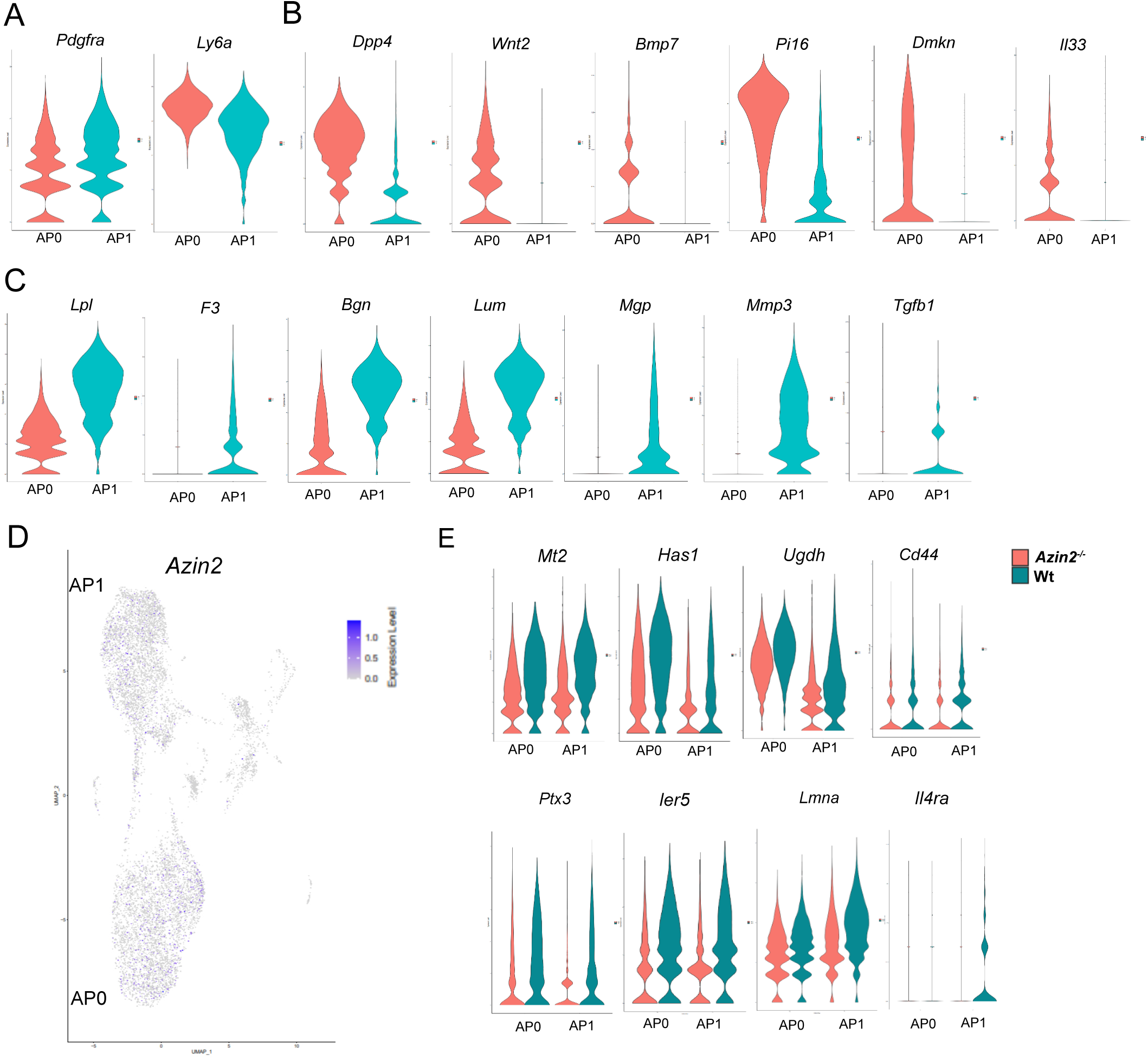
AZIN2 deficiency modulates AP identity, related to Figure 6. scRNA-seq was performed in CD45^-^CD31^-^ SAT SVF cells of wt and *Azin2^-/-^* mice fed for 8 weeks a HFD (2 mice pooled per genotype). **A-C,E.** Violin plots showing gene expression of *Pdgfra* and *Ly6a* (**A**), *Dpp4*, *Wnt2*, *Bmp7*, *Pi16*, *Dmkn*, and *IL33* (**B**) and *Lpl*, *F3* (*Cd142*)*, Bgn*, *Lum*, *Mgp*, *Mmp3*, and *Tgfb1* (**C**) in AP in wt HFD mice, and *Mt2*, *Has1*, *Ugdh*, *Cd44*, *Ptx3*, *Ier5*, *Lmna*, and *Il4ra* in APs of wt and *Azin2^-/-^* HFD mice (**E**). **D.** Two-dimensional UMAP representation of 10,275 cells colored, according to *Azin2* expression (blue) in wt HFD mice.

**Figure S9.**
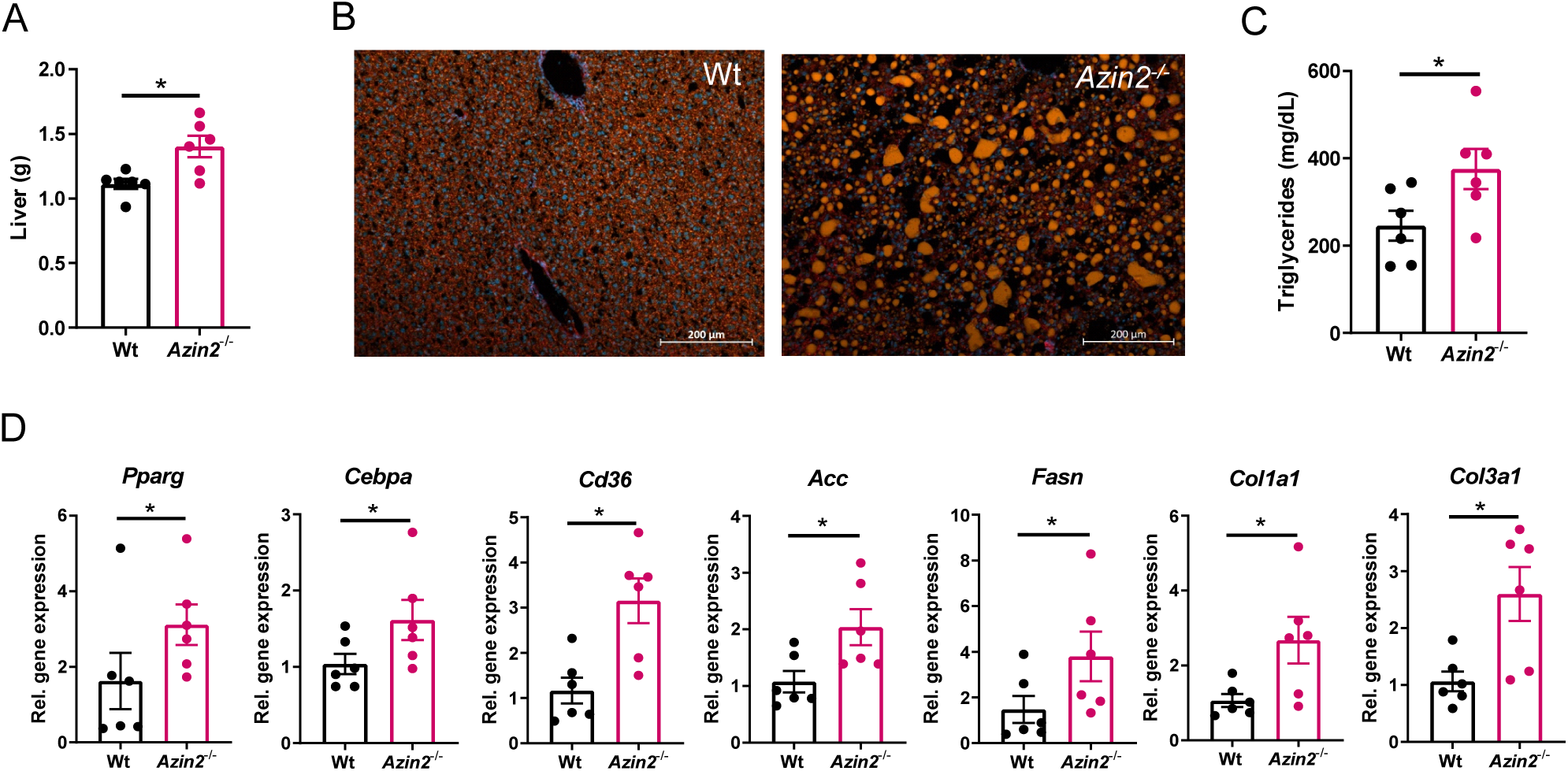
AZIN2 deficiency enhances hepatosteatosis in obesity. Liver weight (**A**), Nile Red staining in liver (**B**), triglyceride content (**C**) and gene expression of *Pparg*, *Cebpa*, *Cd36*, *Acc*, *Fasn*, *Col1a1* and *Col3a1* in liver (**D**) in wt and *Azin2^-/-^* mice were fed for 8 weeks a HFD (n=6 mice per group). Data in **A,C** and **D** are shown as mean±SEM. Mann-Whitney U test was performed *p < 0.01.

**Figure S10.**
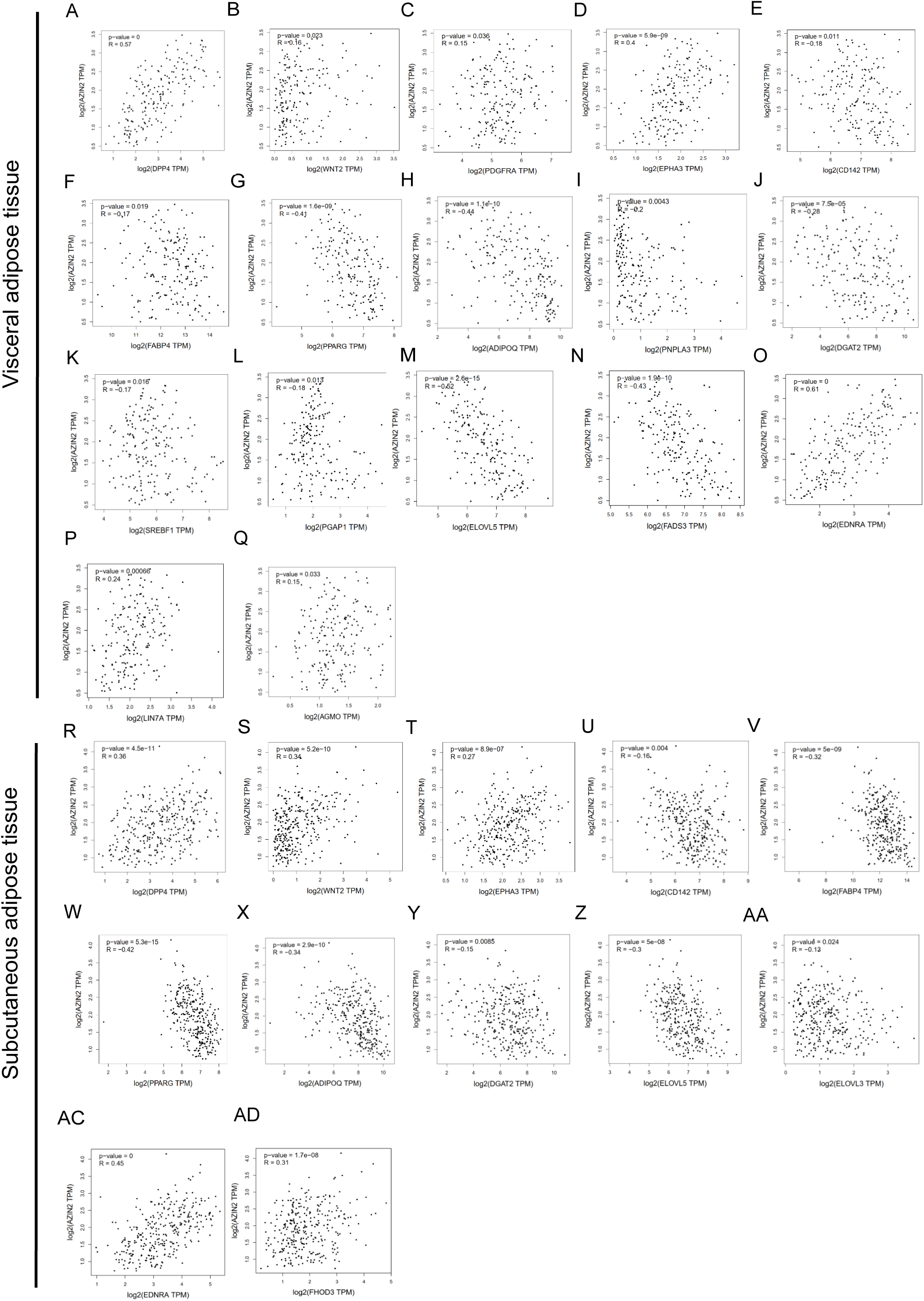
*AZIN2* expression negatively correlates with adipogenesis markers in the human adipose tissue. **A-AD**. Correlation of *AZIN2* expression with *DPP4*, *WNT2*, *PDGFRA*, *EPHA3*, *CD142*, *FABP4*, *PPARG*, *ADIPOQ*, *PNPLA3*, *DGAT2*, *SREBF1*, *PGAP1*, *ELOVL5*, *FADS3*, *EDNRA*, *LIN7A*, and *AGMO* in visceral (A-Q) and *DPP4*, *WNT2*, *EPHA3*, *CD142*, *FABP4*, *PPARG*, *ADIPOQ*, *DGAT2*, *ELOVL5*, *ELOVL3*, *EDNRA*, and *FHOD3* in subcutaneous adipose tissue (R-AD) based on the GEPIA2 database (http://gepia2.cancer-pku.cn/#index).

**Table S1.**
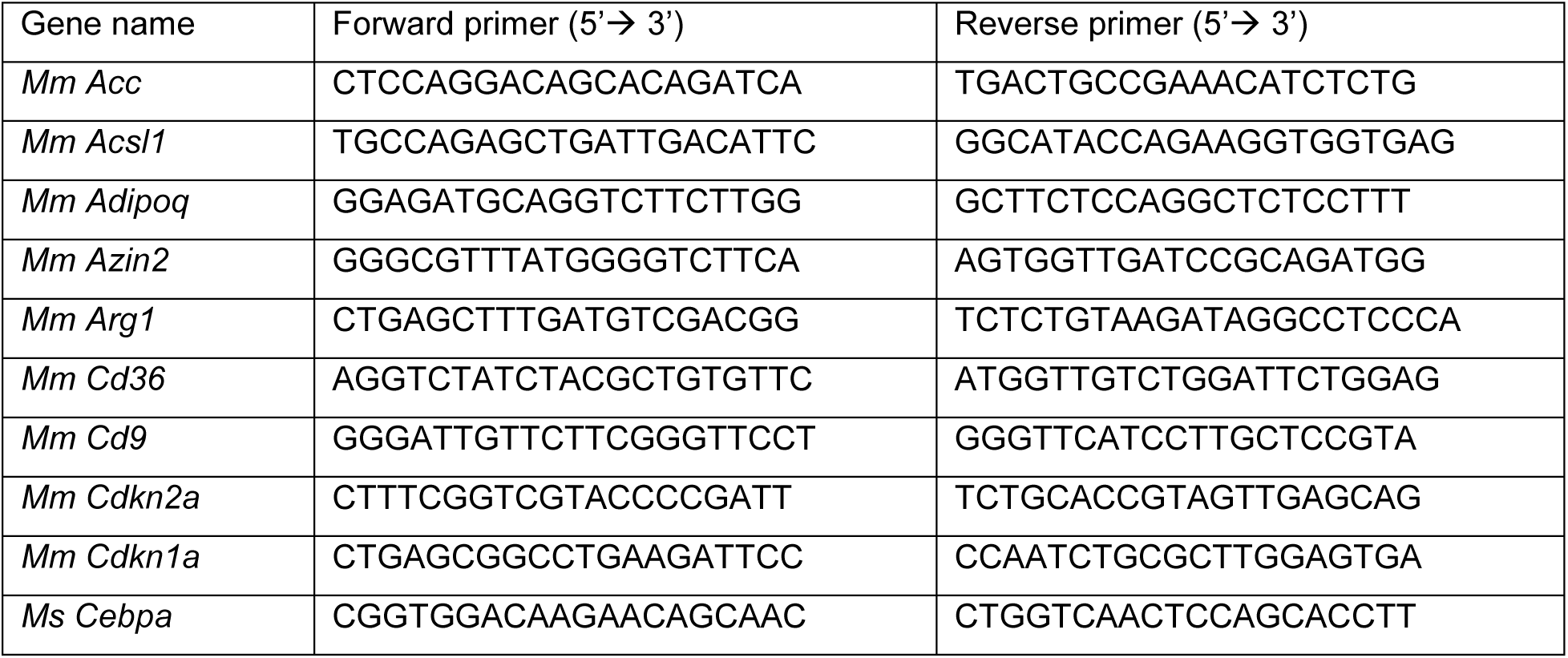
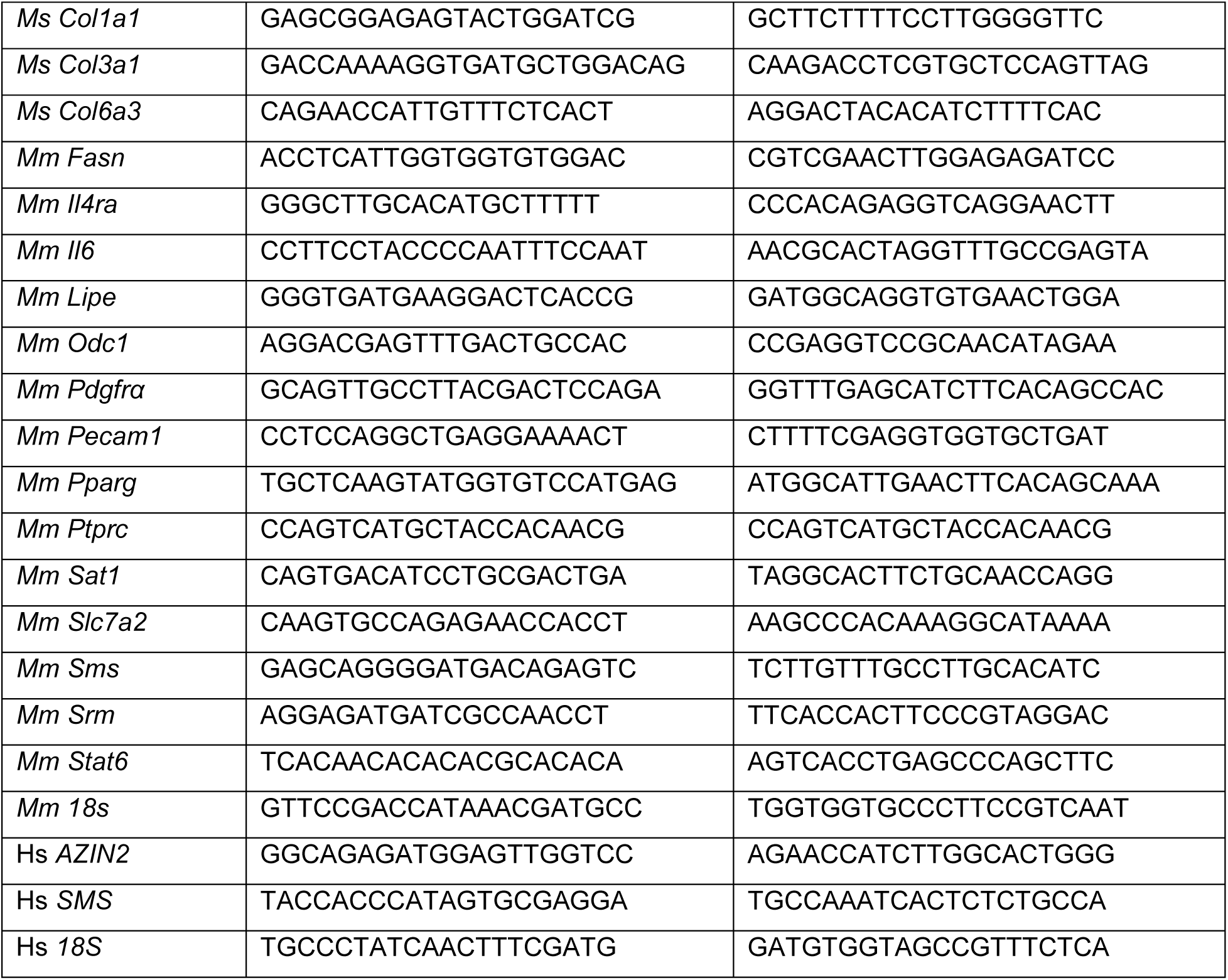

